# Colonial choanoflagellate isolated from Mono Lake harbors a microbiome

**DOI:** 10.1101/2021.03.30.437421

**Authors:** K. H. Hake, P.T. West, K. McDonald, D. Laundon, A. Garcia De Las Bayonas, C. Feng, P. Burkhardt, D.J. Richter, J.F. Banfield, N. King

## Abstract

Choanoflagellates offer key insights into bacterial influences on the origin and early evolution of animals. Here we report the isolation and characterization of a new colonial choanoflagellate species, *Barroeca monosierra,* that, unlike previously characterized species, harbors a microbiome. *B. monosierra* was isolated from Mono Lake, California and forms large spherical colonies that are more than an order of magnitude larger than those formed by the closely related *Salpingoeca rosetta*. By designing fluorescence *in situ* hybridization probes from metagenomic sequences, we found that *B. monosierra* colonies are colonized by members of the halotolerant and closely related *Saccharospirillaceae* and *Oceanospirillaceae,* as well as purple sulfur bacteria (*Ectothiorhodospiraceae*) and non-sulfur *Rhodobacteraceae.* This relatively simple microbiome in a close relative of animals presents a new experimental model for investigating the evolution of stable interactions among eukaryotes and bacteria.

**IMPORTANCE:** The animals and bacteria of Mono Lake (California) have evolved diverse strategies for surviving the hypersaline, alkaline, arsenic-rich environment. We sought to investigate whether the closest living relatives of animals, the choanoflagellates, exist among the relatively limited diversity of organisms in Mono Lake. We repeatedly isolated members of a single species of choanoflagellate, which we have named *Barroeca monosierra,* suggesting that it is a stable and abundant part of the ecosystem. Characterization of *B. monosierra* revealed that it forms large spherical colonies that each contain a microbiome, providing an opportunity to investigate the evolution of stable physical associations between eukaryotes and bacteria.

## DISCOVERY REPORT

### A newly identified choanoflagellate species forms large colonies that contain a microbiome

Choanoflagellates are the closest living relatives of animals and, as such, provide insights into the origin of key features of animals, including animal multicellularity and cell biology [1, 2]. Over a series of four sampling trips to Mono Lake, California (Fig. 1A; Table S1) we collected single-celled choanoflagellates and large spherical choanoflagellate colonies, many of which were hollow (Fig. 1B) and resembled the blastula stage of animal development. In colonies and single cells, each cell had the typical collar complex observed in other choanoflagellates: an apical flagellum surrounded by a collar of microvilli [1, 2]. In these “rosette” colonies, the cells were oriented with the basal pole of each cell pointing inwards and the apical flagellum facing out (Fig. 1B).

**Figure 1.**
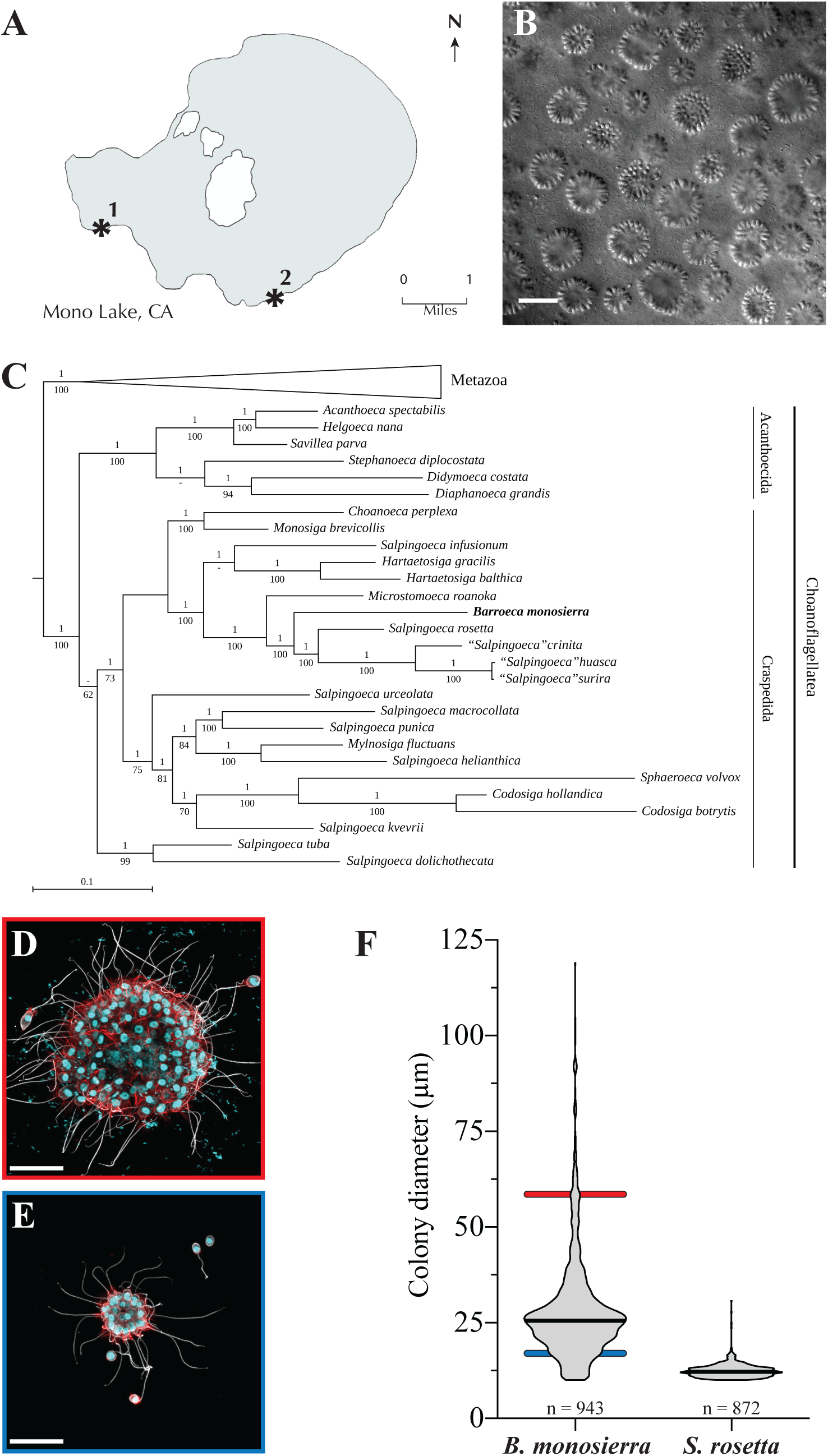
A new rosette-forming choanoflagellate isolated from Mono Lake. (A) Choanoflagellates were collected from two sampling sites (asterisks) near the shore of Mono Lake, California. (Modified from map at monolake.org.) (B) *B. monosierra* forms large rosettes (DIC image). Scale bar = 50µm. (C) *B. monosierra* (shown in bold) is a craspedid choanoflagellate closely related to *S. rosetta* and *Microstomoeca roanoka.* Phylogeny based on sequences of 3 genes: 18S rRNA, EFL and HSP90. Metazoa (seven species) were collapsed to save space. Bayesian posterior probabilities are indicated above each internal branch, and maximum likelihood bootstrap values below. (A ‘-’ value indicates a bifurcation lacking support or not present in one of the two reconstructions.) (D-E) Two representative rosettes reveal the extremes of the *B. monosierra* rosette size range (D, 58 µm diameter; E, 19 µm diameter; scale bar = 20 µm). In *B. monosierra* rosettes, each cell is oriented with its apical flagellum (white; labeled with anti-tubulin antibody) and the apical collar of microvilli (red; stained with phalloidin) pointing out. Nuclei (cyan) were visualized with the DNA-stain Hoechst 33342. (F) Rosettes of *B. monosierra* span from 10 µm in diameter, a size comparable to that of small *S. rosetta* rosettes, to 120 µm, over an order of magnitude larger. Diameters of *B. monosierra* and *S. rosetta* rosettes were plotted as a violin plot; median indicated as thick black line. Diameters of representative rosettes indicated as colored bars behind violin plot (D, red bar; E, blue bar).

To study the Mono Lake choanoflagellates in greater detail, we established clonal strains from ten independent isolates and stored them under liquid nitrogen for future study. Two of the strains were each started from a single-celled choanoflagellate (Fig. S1A, B) and the remaining eight were each started from a single rosette (Table S1). The two strains started from single-celled choanoflagellates, isolates ML1.1 and ML1.2, took on the colonial morphology observed in the other isolates after culturing in the laboratory, suggesting that the colonies and single cells isolated from Mono Lake could belong to the same species. We are aware of no prior reports of choanoflagellates having been cultured from any alkaline soda lake, including Mono Lake.

The 18S rRNA genes for six of the Mono Lake isolates were sequenced and found to be >99% identical (Table S1; Fig. S1C). In further phylogenetic analyses based on 18S rRNA and two protein-coding genes from isolate ML2.1 (Fig. 1C) [3], we found that its closest relatives are the emerging model choanoflagellate *S. rosetta* [4–8], additional *Salpingoeca* spp. [9] and *Microstomoeca roanoka* [3, 10]. The phylogenetic distance separating the Mono Lake species from its closest relatives is similar to the distance separating other choanoflagellate genera. Therefore, we propose the name *Barroeca monosierra*, with the genus name inspired by esteemed choanoflagellate researcher Prof. Barry Leadbeater and the species name inspired by the location of Mono Lake in the Sierra Nevada mountain range. (See Supplemental Methods for further details and a formal species description.)

Although *B. monosierra* and *S. rosetta* form rosette-shaped spherical colonies, they differ greatly in size. *S. rosetta* rosettes range from 10-30 µm in diameter while *B. monosierra* forms among the largest choanoflagellate rosettes observed [1, 11], with a single culture exhibiting rosette sizes spanning from 10-120 µm in diameter (Fig. 1D-F). Unlike the rosettes of *S. rosetta*, in which the basal poles of cells are closely apposed in the rosette center [5,7,11,12], cells in large *B. monosierra* rosettes form a shell on the surface of a seemingly hollow sphere. Inside the ostensibly hollow sphere, in a space equivalent to an epithelial-bound lumen, a branched network of extracellular matrix connects the basal poles of all cells (Fig. S2.)

Upon staining *B. monosierra* with the DNA dye Hoechst 33342, we observed the expected toroidal nuclei in each choanoflagellate cell [12, 13], but were surprised to detect Hoechst-positive material in the interior lumen of *B. monosierra* rosettes (Fig. 2A, A’). Transmission electron microscopy revealed the presence of 1 µm and smaller cells with diverse morphologies bounded by cell walls in the centers of rosettes (Fig. 2B, B’; Fig. S3). Together, these observations led us to hypothesize that the centers of *B. monosierra* rosettes contain bacteria.

**Figure 2.**
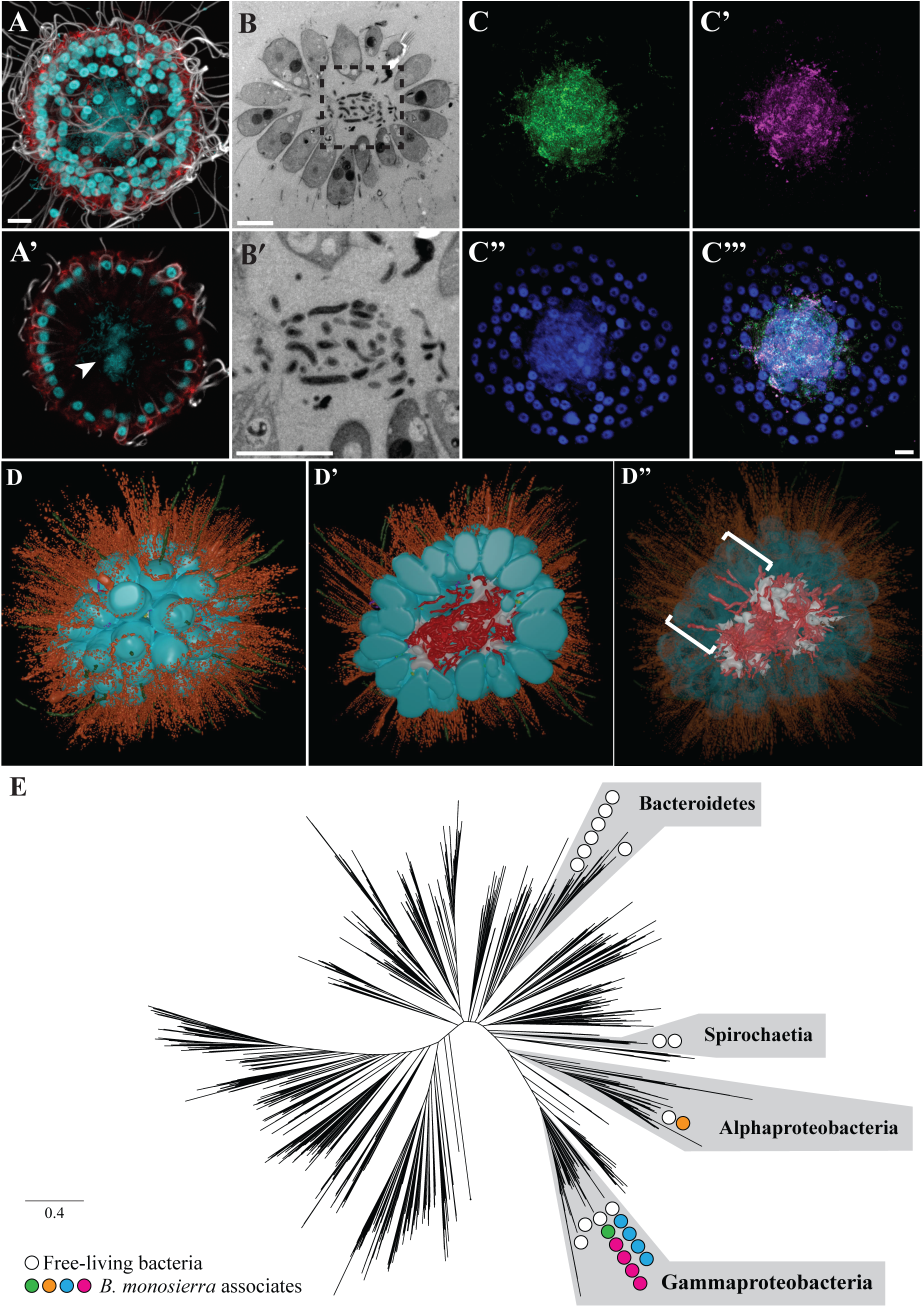
*B. monosierra* rosettes are filled with bacteria. (A, A’) The center of a representative *B. monosierra* rosette, shown as a maximum intensity projection (A) and optical z-section (A’), contains DNA (revealed by Hoechst 33342 staining; cyan). Apical flagella were labeled with anti-tubulin antibody (white); microvilli were stained with phalloidin (red). Hoechst 33342 staining (cyan) revealed the toroidal choanoflagellate nuclei along the rosette perimeter and an amorphous cloud of DNA sitting within the central cavity formed by the monolayer of choanoflagellate cells. (B-B’) Thin section through a representative *B. monosierra* rosette, imaged by transmission electron microscopy (TEM), revealed the presence of small cells in the central cavity. (B’) Inset (box; panel B’) reveals that the interior cells are each surrounded by a cell wall. (C-C’’’) The small cells inside *B. monosierra* rosettes are bacteria, as revealed by hybridization with a broad spectrum 16S rRNA probe (C, green) and a probe targeting Gammaproteobacteria (C’, red). Choanoflagellate nuclei and bacterial nucleoids were revealed by staining with Hoechst (C’’, cyan). (C’’’) Merge of panels C – C’’. Scale bar for all = 5 μm. (D-D’’) 3D reconstruction of a 70-cell *B. monosierra* choanoflagellate rosette from transmission electron micrographs of serial ultrathin sections revealed that the bacteria are closely associated with and wrapped around the ECM inside the rosette. (D) Whole rosette view. (D’) Cut-away view of rosette center. Color code: cell bodies (cyan); microvilli (orange); flagella (green); bacteria (red); ECM (white); intercellular bridges (yellow, see also Fig. S5); filopodia (purple). (D’’) Reducing the opacity of the choanoflagellate cell renderings revealed the presence of bacteria positioned between the lateral surfaces of choanoflagellate cells (brackets, see also Fig. S7). (E) Unrooted phylogenetic tree based on 16 concatenated ribosomal protein sequences representing bacterial diversity modified from [42], illustrated to indicate the phylogenetic placement of bacteria co-cultured from Mono Lake with *B. monosierra.* Scale bar represents the average number of substitutions per site. The bacteria belonged to four major classes: Spirochaetia, Alphaproteobacteria, Gammaproteobacteria, and Bacteroidetes, however the bacteria found associated with *B. monosierra* rosettes came only from Alphaproteobacteria and Gammaproteobacteria. Circles represent the phylogenetic placement of environmental bacteria (white) and choanoflagellate-associated bacteria (*Oceanospirillaceae sp.,* magenta*; Saccharospirillaceae sp.,* green*; Ectothiorhodospiraceae sp.,* blue*; Roseinatronobacter sp.,* orange). See also Figs. S10 and S11. The tree data file is available as Supplemental Data File 1 and can be opened in FigTree or iTOL.

By performing hybridization chain reaction fluorescence *in situ* hybridization (HCR-FISH [14–16]) with a broad-spectrum probe of bacterial 16S rRNA, EUB338 [17], we confirmed that the cells in the central lumen are bacteria (Fig. 2C). A second probe that specifically targeted 16S rRNA sequences from Gammaproteobacteria, GAM42a, revealed that the majority of the bacteria in the centers of rosettes are Gammaproteobacteria (Fig. 2C’)[18]. Finally, by incubating *B. monosierra* cultures with fluorescently labeled D-amino acids, which are specifically incorporated into the cell walls of growing bacteria, we found that the bacteria in *B. monosierra* rosettes are alive and growing (Fig. S4)[19]. Therefore, the microbial community contained within *B. monosierra* represents a microbiome as defined by [20].

To visualize the spatial distribution of choanoflagellate and bacterial cells in a representative rosette, we generated a 3D reconstruction from serial sections imaged by TEM. The rosette contained 70 choanoflagellate cells that were tightly packed, forming a largely continuous monolayer of cells (Fig. 2D). As observed by immunofluorescence microscopy (Figs. 2A and 2A’), all cells were highly polarized and oriented with their apical flagella and collars extending away from the centroid of the rosette. Many cells were connected by fine intercellular bridges (Fig. S5) that have been previously observed in other colonial choanoflagellates, including *S. rosetta* [5, 12].

The 3D reconstruction also revealed at least 200 bacterial cells in the center of the rosette (Fig. 2D’, 2D’’), some of which were physically associated with and wrapped around the choanoflagellate ECM (Fig. S6). A small number of bacterial cells were observed between the lateral surfaces of choanoflagellate cells, although it was not possible to determine whether they were entering or exiting the rosette (Fig. 2D’’; Fig. S7). Rosettes failed to incorporate bacteria-sized bovine serum albumin (BSA)-coated latex microspheres (0.2 µm and 1 µm) into their centers, suggesting that environmental bacteria may not be capable of passively accessing the central lumen of *B. monosierra* colonies (Fig. S8).

### Gammaproteobacteria and Alphaproteobacteria in a *B. monosierra* microbiome

We next sought to identify which bacteria comprise the microbiota of *B. monosierra.* To identify candidate bacteria for which to design FISH probes, we sequenced and assembled metagenomes and 16S rRNA sequences from choanoflagellate-enriched samples and from environmental bacteria-enriched samples. These samples were derived from two co-cultures of *B. monosierra* with Mono Lake bacteria, ML2.1E and ML2.1G (Fig. S9), with the enrichment for choanoflagellates or bacteria performed by centrifugation. A total of 24 different bacterial phylotypes were identified using two complementary bioinformatic approaches (Genome-resolved metagenomic analysis and EMIRGE 16S rRNA Analysis; Supplemental Methods; Table S2 and S3), of which 22 phylotypes were present in fractions enriched with *B. monosierra* rosettes (Table S4). The phylogenetic relationships among these and other bacterial species were determined based on analysis of highly conserved ribosomal proteins and 16S rRNA sequences (Fig. 2E and S10). Not surprisingly, the bacteria maintained in culture with *B. monosierra* represent a subset of the bacterial diversity previously detected in metagenomic analyses of Mono Lake [21].

The 22 bacterial phylotypes detected in cultures with *B. monosierra* may have co-sedimented with the *B. monosierra* rosettes due their community-structure densities (e.g. biofilms), a transient association with the choanoflagellate rosettes (e.g. as prey), or through a stable association with the choanoflagellate rosettes. Upon investigation by FISH microscopy, we detected ten or eleven of these phylotypes in the centers of *B. monosierra* rosettes (Table S4, Fig. S11). (The uncertainty regarding the precise number of choanoflagellate-associated bacterial species stems from the inability to disambiguate 16S rRNA sequences corresponding to one or two of the species.) Of these bacteria, nine were Gammaproteobacteria from the families *Oceanospirillaceae* (Fig. S11A; OceaML1, OceaML2, OceaML3, OceaML4, OceaML4)*, Ectothiorhodospiraceae* (Fig. S11B; EctoML1, EctoML2, EctoML3, EctoML4), and *Saccharospirillaceae* (Fig. S11C; SaccML), matching our original observation that the majority of the bacteria were Gammaproteobacteria (Fig. 2C, C’). The remaining phylotypes was a *Roseinatronobacter* sp. (RoseML; Alphaproteobacteria) (Fig. S11D). The microbiome bacteria exhibited an array of morphologies, from long and filamentous to rod shaped (Figs. S3 and S12). Intriguingly, with the exception of OceaML3, which was exclusively detected inside *B. monosierra* rosettes of ML2.1E (Fig. S13), all other microbiome phylotypes identified in this study were detected both inside and outside the rosettes.

Only one phylotype tested, OceaML1, was found in the microbiome of all *B. monosierra* rosettes (Fig. S14A). The other most frequently observed members of the microbiome were SaccML (93.3% of rosettes), EctoML3 (91.8% of rosettes) and EctoML1 (82.4% of rosettes; Fig. S14A). Two other Gammaproteobacterial phylotypes, OceaML2 and EctoML2, were found in ∼50 - 60% of rosettes, while the Alphaproteobacterium RoseML was found in only 13.9% of rosettes. The most common resident of the *B. monosierra* microbiome, OceaML1, was also the most abundant, representing on average 66.4% of the total bacterial load per rosette (Fig. S14B). Other abundant bacteria, some found in >80% of rosettes, represented rather smaller percentages of the average bacterial biomass in the microbiomes in which they were found. For example, SaccML was found in 93.3% of microbiomes but represented only 30.3% of total bacteria in the *B. monosierra* rosettes in which it was found. EctoML1, which was found in 82.4% of *B. monosierra* rosettes represented less than 10% of the bacteria in the bacteria in which was detected.

## Discussion

Interactions with bacteria are essential to choanoflagellate nutrition and life history. Bacteria are the primary food source for choanoflagellates, and the choanoflagellate *S. rosetta* responds to different secreted bacterial cues to undergo either multicellular developmental or mating [26–29]. We report here the isolation and characterization of a new choanoflagellate species, *B. monosierra,* that forms large rosettes and contains a microbiome. To our knowledge, this is the first example of a stable physical interaction between choanoflagellates and bacteria. Future studies will be important to determine how the *B. monosierra* rosettes and their bacterial residents interact.

*B. monosierra* and its associated microbiome provide a unique opportunity to characterize the interaction between a single choanoflagellate species and a microbial community. It is important to note that the host-microbe associations reported here were identified in lab-grown *B. monosierra* and it will be necessary in the future to investigate the composition of the microbiome in wild populations from Mono Lake. Due to the phylogenetic relevance of choanoflagellates, the relationship between *B. monosierra* and its microbiota has the potential to illuminate the ancestry of and mechanisms underlying stable associations between animals and bacteria, colonization of eukaryotes by diverse microbial communities, and insights into one of the most complex animal-bacterial interactions, the animal gut microbiome.

## Data Availability

Genbank accession numbers for bacterial 16S rRNA sequences are listed in Table S5. Sequences for *B. monosierra* 18S rRNA, EFL, and Hsp90 (Fig. 1C) have been assigned GenBank accession numbers MW838180, MW979373 and MW979374, respectively. 18S sequences for different *B. monosierra* strains (Fig. S1, Table S1) have been assigned GenBank accession numbers MZ015010-MZ015015. The assembled *B. monosierra* genome sequence is currently being uploaded to GenBank and will be available under the accession number PRJNA734368. The *B. monosierra* genome, all bacterial genome sequences, and all relevant input and output data from the phylogenetic trees presented in Fig. 1C and Fig. S1 are available via FigShare (https://doi.org/10.6084/m9.figshare.14474214).

## Author contributions

P.W. performed metagenomic analysis, K.M. did the transmission electron microscopy, D.L. and P.B. did the TEM-reconstruction, and C.F. helped culture B. monosierra. A.G.D.L.B. imaged the theca. K.H. performed all other experiments and analysis. D.J.R. originally isolated B. monosierra and contributed to manuscript editing and the taxonomic description. J.B. and N.K. contributed to project leadership, experimental design, figure design, writing, and editing.

## Acknowledgements

We thank Tarja Hoffmeyer for help with establishing some of the original *B. monosierra* cultures. We thank Emil Ruff and Julia Schwartzman for assistance with early CARD-FISH experiments through the Physiology Course and Microbial Diversity Course at the Marine Biological Laboratory in Woods Hole. We thank Michael VanNieuwenhze for the fluorescent D-amino acids. Frank Nitsche provided advice on the formulation of an artificial Mono Lake culture medium. We thank the following for helpful discussions and research support: Reef Aldayafleh, Cédric Berney, David Booth, Ben Larson, Monika Sigg, Laura Wetzel, and Arielle Woznica. We thank Karen Carniol for valuable feedback on the manuscript. This material is based upon work supported by the National Science Foundation Graduate Research Fellowship under Grant No. DGE 1106400 and DGE 1752814. D.J.R. received the support of a fellowship from ”la Caixa” Foundation (ID 100010434) with the fellowship code LCF/BQ/PI19/11690008.

## SUPPLEMENTARY MATERIALS

### Text S1: Supplemental Materials and Methods

**We propose the creation of a new genus for the Mono Lake species for the following four reasons:**

1. The phylogenetic distance separating the Mono Lake species from its closest relatives (*M. roanoka* and *S. rosetta*) is equivalent to the distance between genera in other parts of the choanoflagellate tree (Fig. 1C; for example, between *C. perplexa* and *M. brevicollis*).
2. The internal branch joining the Mono Lake species together with the *S. rosetta* group is short, indicating that they probably did not diverge long after the split from *M. roanoka*. This is similar to the situation observed within the Acanthoecida (where short internal branches lead to many separate genera) and stands in contrast to other bona fide genera in Craspedida that do share a long internal branch, such as *Hartaetosiga* or *Codosiga*.
3. It would be preferable to avoid adding another species to the *Salpingoeca* genus, which is already highly paraphyletic. Adding another *Salpingoeca* would increase confusion when interpreting the relationship among species (which outweigh the disadvantages of creating a monospecific genus, of which numerous examples currently exist within Craspedida: for example, *Microstomoeca* and *Mylnosiga*). Furthermore, as the type species of *Salpingoeca* has not yet been sequenced, creating a new genus for the Mono Lake species would avoid the possibility of its needing to be transferred to another genus at a later date.
4. Choanoflagellates and animals diverged at least 600 million years ago, and the average phylogenetic distance between any two choanoflagellates is at least as great as the average phylogenetic distance between any two animals [1]. Thus, the Mono Lake species is likely to have experienced at least tens of millions of years of independent evolution after separating from its closest known relatives in the tree. In fact, a recent study, which did not include the Mono Lake species, estimated the divergence of *M. roanoka* and *S. rosetta* to have occurred roughly 100 million years ago, and the divergence of *S. rosetta* from its closest relatives to have occurred roughly 38 million years ago [2].

#### Taxonomic Summary

Order Craspedida Cavalier-Smith 1997 [3]

Family Salpingoecidae Kent (1880–1882), emend. sensu Nitsche et al. 2011 [4]

*Barroeca* gen. nov. Hake, Burkhardt, Richter and King

Uninucleated microbial eukaryote with a single, centrally positioned apical flagellum, which is surrounded by a collar of actin-supported microvilli. Phagotrophic. At least some species possess an organic theca. Phylogenetically more closely related to *Barroeca monosierra* than to *Microstomoeca roanoka* or *Salpingoeca rosetta*.

**Etymology**: The genus is named for Barry S. C. Leadbeater, the author of numerous research articles and the definitive book on choanoflagellates, and a remarkably positive influence on choanoflagellate research and researchers throughout a career spanning more than 50 years.

**Type species:** *Barroeca monosierra* Hake, Burkhardt, Richter and King

*Barroeca monosierra* Hake, Burkhardt, Richter and King

**Etymology**: *mono* was derived from the source locality, Mono Lake, and *sierra* for the Sierra Nevada mountain range in which Mono Lake is found.

**Type locality**: Shore of Mono Lake, California (37°58’42.7”N 119°01’52.9”W).

**Description**: Cell body is ∼6-7 μm long. Apical microvillous collar is ∼7.5-9.5 μm long. Apical flagellum is ∼20-25 μm long. Single cells may be found attached to a substrate via a long (∼30 μm) basal pedicel (Fig. S1). Single cells may possess an organic cup-shaped theca with a distinctive ∼0.75 μm outward-facing lip on its apical end (Fig. S1). Rosette colonies can be as large as 125 μm in diameter and consist of a spheroidal arrangement of cells surrounding a hollow space containing microbiome bacteria. Adjacent cells connect via intercellular bridges, surrounded by shared plasma membrane and positioned slightly basal to cell equators. Intercellular bridges are cylindrical structures, 200-300 nm wide and 300-500 nm long, partitioned by two parallel densely osmophilic plates 175-275 nm apart.

**Type material:** The strain ML 2.1 is the one used for describing this species and is illustrated in Figures 1 and 2.

**Gene sequence:** The partial small subunit ribosomal RNA (SSU rRNA) gene sequence of strain ML2.1 has been deposited in GenBank, accession code MW838180.

#### Initial isolation of choanoflagellate *B. monosierra*

*B. monosierra* was originally isolated from water samples collected at Mono Lake, CA from 2012-2014 (Table S1; Fig. 1A). 30 mL of lake water was collected in a T25 cell culture flask with a vented cap (Thermo Fisher Scientific, Waltham, MA; Cat. No. 10-126-28). The flasks were placed in the dark for 2-4 weeks at 22°C to reduce the load of photosynthetic microorganisms and allow the choanoflagellates to grow and increase in density. Flasks were visually screened for the presence of choanoflagellate cells, and the choanoflagellates were clonally isolated by two serial dilution-to-extinction steps, similar to the method described in (Fig. S9)[5]

#### Choanoflagellate Growth Media

*B. monosierra* was initially grown in 0.22 µm filtered Mono Lake water collected from the same location as the isolation (Table S1) and later transferred into artificial Mono Lake water (AFML) designed to approximate the water chemistry of Mono Lake, based on assessments by Dr. Frank Nitsche (University of Cologne) and the Los Angeles Department of Water and Power (LADWP; http://www.monobasinresearch.org/images/mbeir/dchapter3/table3b-2.pdf). AFML was prepared by adding salts and minerals (Table S7) in order to MilliQ water and filtering through a 0.22 µm filter without autoclaving. Calcium chloride dihydrate was added as a stock solution (1:1000). 1000x Trace elements and 1000x L1 Vitamins were added to AFML right before use as described in [6] where they were added to artificial sea water (ASW) for culturing *S. rosetta*. Mono Lake Cereal grass medium (CGM3ML) was made by infusing unenriched freshly autoclaved AFML with cereal grass pellets (5g/L) (Carolina Biological Supply, Burlington, NC; Cat. No. 132375) as previously described [7]. Davis Mingolis synthetic media (DMSM)[8] was made by adding the chemicals described here (Table S7) to MilliQ water and sterile filtering through a 0.22 µm filter without autoclaving. *S. rosetta* was propagated in unenriched sea water (32.9 g Tropic Marin sea salts to 1L water) with 5% (vol/vol) sea water complete medium [9] resulting in HN medium (250 mg/L peptone, 150 mg/L yeast extract, and 150 µl/L glycerol in unenriched sea water) as previously described [7].

#### Culturing conditions and establishment of ML2.1 Cultures

Clonal isolates of *B. monosierra* cultures were passaged once a week by adding 3 mL of culture to 9 mL of 10% CGM3ML diluted in AFML with added vitamins and trace minerals into a T75 vented cell-culture flask (Fig. S9). Frozen stocks were prepared as previously described [10]. The optimal growth conditions for *B. monosierra* were tested by growing the culture in different concentrations of media previously used for other choanoflagellate species [1,7,11] as well as media used to isolate bacteria from alkaline soda lakes [8]. We found that *B. monosierra* grew best under two conditions (Fig. S9, Box 2). The first condition involved passaging 3 mL of *B. monosierra* culture in 9 mL of 2.5% DMSM diluted in AFML and resulted in the culture ML2.1E. The second involved passaging 3 mL of *B. monosierra* in 9 mL of 10% CGM3ML diluted in AFML that was treated with Gentamicin (50 µg/mL) over six weeks and then maintained in 10% CGM3ML diluted in AFML resulting in the culture ML2.1G.

Cultures ML2.1E and ML2.1G were passaged under these conditions for 4 months before being sequenced (Fig. S9, Box 3). Shortly after sequencing, ML2.1G became overpopulated with bacteria that outcompeted the choanoflagellates. We were not able to recover this culture from a freeze down. Due to the variability in growth of ML2.1E, likely due to the diversity of bacterial prey and their growth dynamics, we found it was helpful to always have two cultures growing in different media to improve the chances of having a healthy culture on hand. Therefore, we began passaging ML2.1E in not only 2.5% DMSM, but also 10% CGM3ML resulting in the culture ML2.1EC (Fig. S9, Box 4). Experiments were performed in either ML2.1E or ML2.1EC based on the quality of the cultures on the day of the experiment.

#### High-throughput colony size analysis

For the colony size analysis in Fig. 1F, the ML2.1G culture was used for the *B. monosierra* sample. For *S. rosetta,* the colony strain Px1 (ATCC PRA-366: https://www.atcc.org/Products/All/PRA-366.aspx), a monoxenic culture consisting of *S. rosetta* and the rosette-inducing bacterium *A. machipongonensis,* was used for analysis [12]. Rosettes were concentrated ten-fold by centrifugation at 2000xg for 5 min. Cells were stained with LysoTracker Red DND-99 (Thermo Fisher Scientific; Cat. No. L7528) at a concentration of 1 µl per 100 µl of cells. This concentration of LysoTracker selectively labels the entire cell body of the choanoflagellate and not prey bacteria.

Colonies were imaged by immediately placing a 20 µl drop of stained cells on a glass slide and gently squishing with a 24×40 coverslip (Fisher Scientific; Cat. No. 12-545D). Slides were imaged immediately using a Zeiss Axio Observer.Z1/7 Widefield microscope with a Hamamatsu Orca-Flash 4.0 LT CMOS Digital Camera (Hamamatsu Photonics, Hamamatsu City, Japan) and 20X/NA 0.8 Plan-Apochromat (Zeiss). Four 100-image tilescans with 0% image overlap were acquired resulting in 400 images per sample. (Fig. 1D and E).

Images were analyzed on FIJI [13] and scanned for quality by hand. Any image with debris or out of focus choanoflagellate colonies was deleted. An automatic threshold was set using the intermodes method [14]. Measurements were set to collect area and Feret’s diameter. Images were analyzed using the ‘Analyze Particles’ command with settings to exclude on edges and to include holes. Minimum rosette area and diameter were set to 35 µm and 10 µm, respectively, based on *S. rosetta* measurements of 2-cell rosettes. Total number of rosettes imaged were 943 for *B. monosierra,* and 872 for *S. rosetta.* Data was presented as a violin boxplot made using GraphPad Prism8 showing the median area or diameter (black line), and the kernel density trace (black outline) plotted symmetrically to show frequency distribution for the given measurements.

#### Immunofluorescence, confocal imaging, and live cell microscopy

For immunofluorescence staining of *B. monosierra* (Fig. 1D and E; Fig. 2A and A’), a round poly-L-lysine coated coverslip (BD Biosciences) was placed at the bottom of a 24 well plate. 1750 µl of *B. monosierra* culture was added to each well with a coverslip followed by 250 µl of 32% Paraformaldehyde (PFA) resulting in a final concentration of 4%PFA. The 24 well plate was spun at 1000xg for 15 minutes at room temperature to concentrate choanoflagellate colonies onto the coverslip. The fixative and culture media was replaced with PEM (100 mM PIPES-KOH, pH 6.95; 2 mM EGTA; 1 mM MgCl_2_). Immunofluorescence was continued as previously described (13) with 50 ng/ml mouse E7 anti-β-tubulin antibody (Developmental Studies Hybridoma Bank, University of Iowa, Iowa City, IA; Cat. No. AB_2315513), 8 ng/ml Alexa Fluor Plus 488 Goat anti-Mouse IgG (H+L) secondary antibody (Thermo Fisher Scientific; Cat. No. A32723), 4 U/ml Rhodamine phalloidin (Thermo Fisher Scientific; Cat. No. R415), and 0.1mg/mL Hoechst 33342 (Thermo Fisher Scientific; Cat. No. H3570). Coverslips were mounted in Pro-Long Diamond antifade reagent (Thermo Fisher Scientific; Cat. No. P36970) and left to cure overnight before imaging.

Confocal images were acquired using a 63X/NA 1.40 Plan-Apochromatic oil immersion objective on either a Zeiss LSM 700 confocal microscope or a Zeiss LSM 880 Airyscan confocal microscope with an Airyscan detector (Carl Zeiss AG, Oberkochen, Germany). Images acquired using the Airyscan detector were processed using the automated Airyscan algorithm (Zeiss) and then reprocessed with the Airyscan threshold 0.5 units higher than the automated reported threshold.

For live epifluorescence and differential interference contrast (DIC) imaging (Fig. 1B), cultures of either *B. monosierra* or *S. rosetta* were typically concentrated tenfold, stained with a combination of dyes, and imaged similarly to the colony size analysis by placing 20 µl of cell culture on a slide and gently squishing with a 24×40 coverslip. To stain the DNA, cultures were stained ten minutes with 0.1 mg/mL Hoechst 33342 (Thermo Fisher Scientific; Cat. No. H3570). To label the ECM of *B. monosierra* (Fig. S2), fluorescein-labeled Concanavalin A (Con A) (Vector Labs, Burlingame, CA; Cat No. FL-1001), or for *S. rosetta,* fluorescein labeled Jacalin (Vector Labs; Cat No. FL-1151) was added to cultures at a concentration of 5 µg/mL for five minutes. LysoTracker DND-99 (Thermo Fisher Scientific; Cat. No. L7528) labeled the choanoflagellate colonies as described in the colony size analysis. Cultures were imaged without washing the dyes off. Slides were imaged using a Zeiss Axio Observer.Z1/7 Widefield microscope with Hamamatsu Orca-Flash 4.0 LT CMOS Digital Camera and a 100x NA 1.40 Plan-Apochromatic oil immersion objective (Zeiss).

#### Labeling cultures with D-amino acids, and incubating with fluorescent beads

To label growing bacteria with fluorescently labeled D-amino acids (Fig. S4), cultures were concentrated fivefold and HADA D-amino acids were added at a concentration of 2 mM [15] and incubated for 24 hours. The cultures were imaged as described in live cell microscopy. To test rosette permeability (Fig. S8), 2 mL of ML 2.1 grown in AFML was concentrated twofold and 0.2 µm Fluospheres® (Thermo Fisher Scientific; Cat. No. F8848) and 1 µm Fluospheres® (Thermo Fisher Scientific; Cat. No. F8851) were added at a concentration of 1:100. Cultures were left with beads for 24 hours and imaged as described in live cell microscopy.

#### Transmission electron microscopy (TEM) and 3D reconstruction

For the TEM images (Fig. 2B, B’; Fig. S3; Fig. S5-7) we modified methods previously established for *S. rosetta* [16, 17]. We first concentrated 40 mL of cultured *B. monosierra* rosettes by gentle centrifugation (200xg for 30min) resuspended in 20% BSA (Bovine Serum Albumin, Sigma) made up in artificial seawater medium, and concentrated again. Most of the supernatant was removed and the concentrated cells transferred to high-pressure freezing planchettes varying in depth between 50 and 200 μm (Wohlwend Engineering). Freezing was done in a Bal-Tec HPM-010 high-pressure freezer (Bal-Tec AG).

The frozen cells were stored in liquid nitrogen until needed, and then transferred to cryovials containing 1.5 mL of fixative consisting of 1% osmium tetroxide plus 0.1% uranyl acetate in acetone at liquid nitrogen temperature (−195°C) and processed for freeze substitution according to the method described here [18, 19]. Briefly, the cryovials containing fixative and cells were transferred to a cooled metal block at −195°C (the cold block was put into an insulated container such that the vials were horizontally oriented) and shaken on an orbital shaker operating at 125 rpm. After 3 h, the block/cells had warmed to 20°C and were ready for resin infiltration.

Resin infiltration was accomplished according to the method of described here [18]. Briefly, cells were rinsed three times in pure acetone and infiltrated with Epon-Araldite resin in increasing increments of 25% over 30 min plus three changes of pure resin at 10 min each. Cells were removed from the planchettes at the beginning of the infiltration series and spun down at 6,000x g for 1 min between solution changes. The cells in pure resin were placed in between two PTFE-coated microscope slides and polymerized over 2 h in an oven set to 100°C. Images of cells were taken on an FEI Tecnai 12 electron microscope.

For the 3D reconstruction in Fig. 2D-D’’, we collected images from serial sections followed the approach previously described in [17]. Briefly, the cells were cut from the thin layer of polymerized resin and remounted on blank resin blocks for sectioning. Serial sections of varying thicknesses between 70–150 nm were cut on a Reichert-Jung Ultracut E microtome and picked up on 1 x 2-mm slot grids covered with a 0.6% Formvar film. Sections were post-stained with 1% aqueous uranyl acetate for 7 min and lead citrate for 4 min. Images of sections were collected on an FEI Tecnai 12 electron microscope.

The 3D reconstruction of an entire rosette was achieved using serial ultrathin TEM sectioning (ssTEM). 176 images taken from consecutive 150 nm thick sections were compiled into a single stack and imported into the Fiji [20] plugin TrakEM2 [21]. Stack slices were automatically aligned using default parameters with minor modifications (steps per octave scale were increased to 5 and maximal alignment error reduced to 50 px). Automatic alignments were curated and manually corrected if unsatisfactory. Cellular structures were segmented manually, and 3D reconstructed by automatically merging traced features along the z-axis. Meshes were then smoothed in TrakEM2, exported into the open-source 3D software Blender 2.77, and rendered for presentation purposes only.

#### Genomic DNA extraction and sequencing

Colonies and free-living bacteria were separated by differential centrifugation. To enrich for large rosettes, 10 mL of dense culture was concentrated in a clinical centrifuge at 200xg for 30min. The supernatant was transferred to a clean 15 mL conical tube and set aside for bacterial enrichment. We resuspended the rosette pellet twice in 15 mL of AFML and centrifuged at 2000xg for 10 minutes to separate the choanoflagellates from free-living bacteria. The rosette pellet was transferred in residual media to a 1.5 mL tube, confirmed by microscopy that it had enriched, and then was pelleted and frozen under liquid nitrogen. Free-living bacteria were enriched from the first supernatant by centrifuging at 500xg for 10 minutes to deplete single cell choanoflagellates and small rosettes that didn’t pellet in the first spin. The supernatant was concentrated at 4000xg for 20 min. Concentrated bacteria were re-suspended in 1 mL of residual media and filtered through a 3 µm filter to remove single cell choanoflagellates. Bacteria were transferred to a 1.5 mL tube, confirmed by microscopy that there were few choanoflagellates, pelleted, and frozen under liquid nitrogen. A phenol-chloroform extraction was performed to generate gDNA. Samples were sequenced with 150 bp Illumina paired end reads to a depth ranging from 22.4 to 34.1 Gbp.

#### Metagenome assembly, annotation, and binning

Sequencing reads were processed with bbtools (http://jgi.doe.gov/data-and-tools/bbtools/) to remove Illumina adaptors as well as phiX Illumina trace contaminants. Reads were then quality-filtered with SICKLE (https://github.com/najoshi/sickle). IBDA_UD [22] was used to assemble and scaffold filtered reads from each sample with default parameters. Putative eukaryotic scaffolds were filtered with EukRep [23] prior to bacterial binning. Protein coding sequences were predicted using MetaProdigal [24] on whole assembled metagenomic samples. Bacterial 16S sequences were reconstructed with EMIRGE (v. 0.61.0)[25]. The exact number of 16S rRNA sequences present in each sample could not be determined due to the co-assembly of 16S rRNAs from closely related species in shotgun metagenome samples [25]. In addition, 16S sequences frequently do not bin into genomes due to their high copy number, so they could not be tied directly to binned genomes. The choanoflagellate 18S rRNA sequence was identified with an HMM-based approach (https://github.com/christophertbrown/bioscripts), where a bacterial 16S rRNA, archaeal 16S rRNA, and eukaryotic 18s rRNA model were run concurrently and overlapping predictions were picked based on the best alignment. Genome bins were identified and refined using ggKbase (ggkbase.berkeley.edu) to manually check the GC, coverage, and phylogenetic profiles of each bin. dRep [26] was used to de-replicate genomic bins across samples. The results and analyses from each of the metagenome sequencing and assembly processes is summarized in Table S2 and the bacterial community overlap between different samples is displayed in Fig. S15.

#### 18S rRNA sequencing of Mono Lake isolates

We amplified the 18S rRNA gene from 6 Mono Lake choanoflagellate isolates via PCR of genomic DNA with the universal eukaryotic 18S primers 1F (5’ AACCTGGTTGATCCTGCCAGT 3’) and 1528R (5’ TGATCCTTCTGCAGGTTCACC 3’)[27]. We cloned PCR products using the TOPO TA Cloning vector from Invitrogen, following the manufacturer’s protocol. We next performed Sanger sequencing on multiple clones per isolate, using the T7 forward primer and the M13 reverse primer on the vector backbone, which produced between 2-9 successful clones per isolate, after removing empty vector and contaminant sequences. For each isolate, we aligned sequences using FSA version 1.15.7 [28] with the ‘--fast’ option and then inferred a majority-rule consensus sequence from the alignment.

#### Phylogenetic analysis of choanoflagellate sequences

To place the newly isolated Mono Lake species in a phylogenetic context (Figs. 1C and S1), we began with the sequences used for tree reconstruction in [1], selecting only the choanoflagellate species and 7 representative animals (Porifera: *Amphimedon queenslandica*, Ctenophora: *Mnemiopsis leidyi*, Placozoa: *Trichoplax adhaerens*, Cnidaria: *Nematostella vectensis*, Ecdysozoa: *Daphnia pulex*, Deuterostomia: *Mus musculus*, Lophotrochozoa: *Capitella teleta*). We then added the 18S rRNA sequences of three species closely related to *S. rosetta*: *S. crinita*, *S. huasca* and *S. surira* [29]. We built one tree (Fig. S1) incorporating the six 18S sequences we obtained from colonial Mono Lake cultures via PCR (Table S1). Next, we searched the genome sequence for the ML 2.1 isolate for an additional 5 genes used for choanoflagellate phylogeny reconstruction in [30]. We were able to retrieve the sequences of two genes, EFL and HSP90, via BLAST using the *S. rosetta* and *M. brevicollis* genes as queries and taking the top hit (which was the same for both query species in both cases). We incorporated these two genes into a concatenated three-gene phylogeny (Fig. 1C).

Prior to tree reconstruction, we processed each gene independently, as follows. For protein-coding genes (EFL and HSP90), we trimmed poly-A tails for all sequences using the program trimest from the EMBOSS package version 6.6.0.0 [31], with the ‘- nofiveprime’ option and all other parameters left at their defaults. We aligned each gene separately using FSA version 1.15.7 [28] with the ‘--fast’ option, and trimmed the resulting alignments using trimAl version 1.2rev59 [32] with the ‘-gt 0.3’ option.

We concatenated trimmed sequences into a single alignment with three total partitions for tree reconstruction (following [30], a thorough exploration of choanoflagellate phylogenetics, which included the genes and choanoflagellate species we analyzed here, with the exception of *B. monosierra* and the three *S. rosetta* relatives that we added as described above): one partition for the 18S gene, one partition for the first and second codon positions in the protein-coding genes, and one partition for the third codon position. We reconstructed maximum likelihood phylogenies using RAxML version 8.2.4 [33], with the GTRCAT model and the ‘-f a -N 100’ options for bootstrapping. We reconstructed Bayesian phylogenies with MrBayes version 3.2.6 [34], using a GTR + I + Γ model run for 1 million generations, with all other parameter values left at their defaults. The models chosen were according to the precedent set in reference [30]. For the phylogeny with different isolate 18S sequences, the final average standard deviation of split frequencies was 0.004327, and for the three-gene phylogeny for ML 2.1, it was 0.000074.

#### Phylogenetic analysis of bacteria

For generating the bacterial 16 ribosomal protein tree (Fig. 2E), a previously developed data set [35] was used. For each Mono Lake bacterial genome, 16 ribosomal proteins (L2, L3, L4, L5, L6, L14, L15, L16, L18, L22, L24, S3, S8, S10, S17, and S19) were identified by BLASTing [36] a reference set of 16 ribosomal proteins against the protein sets. (Dereplicated bin set used for constructing the ribosomal protein tree is summarized in Table S8.) BLAST hits were filtered to a minimum e-value of 1.0 × 10^−5^ and minimum target coverage of 25%. The resulting 16 ribosomal protein sets from each Mono Lake bacterial genome and the full reference set were aligned with MUSCLE (v. 3.8.31)[37]. Alignments were trimmed by removing columns containing 90% or greater gaps and then concatenated. A maximum likelihood tree was constructed using RAxML (v. 8.2.10)[33] on the CIPRES web server [38] with the LG plus gamma model of evolution (PROTGAMMALG) and the number of bootstraps automatically determined with the MRE-based bootstopping criterion.

For the bacterial 16S rRNA tree (Fig. S10), Mono Lake 16S rRNAs were aligned against the SILVA 128 SSU Ref NR 99 database [39] with BLAST [36] and the top three hits for each individual sequence, as well as a set of archaeal sequences for use as an outgroup, were aligned with MUSCLE (v. 3.8.31)[37][35][34][32]. For both the bacterial 16S rRNA trees, RAxML was used to construct a maximum likelihood tree with the GTRCAT model and the MRE-based bootstopping criterion.

#### Fluorescence *In Situ* Hybridization (FISH) Probe Design

Starting with the metagenome sequences described above, we developed FISH probes directed against the 16S sequence for each of the potential species detected in each sample. A detailed protocol for probe design using the ARB project software (http://www.arb-home.de/)[40] can be found at protocols.io at the following link: https://www.protocols.io/view/16s-rrna-probe-design-for-hcr-fish-wdffa3n. In short,16S rRNA sequences assembled from metagenomic sequencing utilizing EMIRGE above were imported and aligned to the SILVA 128 SSU Ref NR 99 database [39] in the ARB project using the automatic alignment tool. Probes were designed against individual bacteria identified in the *B. monosierra* samples containing choanoflagellate colonies (Fig. S9, Table S4) using the probe design tool and checked with the probe match tool in the ARB project software. The following HCR-amplification sequences were added to the 3’ end of the probe sequences based on the fluorophore (488, 594, 647) and hairpins (B1, B2, B3) intended for the experiment: B1(488)(5’-ATATA GCATTCTTTCTTGAGGAGGGCAGCAAACGGGAAGAG-3’), B2(594)(5’-AAAAA AGCTCAGTCCAT CCTCGTAAATCCTCATCAATCATC-3), and B3(647)( 5’-TAAAA AAAGTCTAATCCGTCCCTGCCTCTATATCTCCACTC-3’). Full length sequences of the probe with spacer and initiator sequences can be found in Table S6.

#### Fluorescence *In Situ* Hybridization and imaging

For the FISH experiments (Fig. 2C-C’’’; Fig. S11-S13) we used hybridized chain reaction (HCR) – FISH. Our detailed protocol for HCR–FISH for choanoflagellate cultures can be found at protocls.io at the following link: https://www.protocols.io/edit/hcr-fish-for-choanoflagellate-cultures-wddfa26/steps. In short, the hairpin solutions and amplifier sequences used in this study were obtained from Molecular Instruments (www.molecularinstruments.com). *B. monosierra* choanoflagellate cultures with free-living bacteria were fixed overnight in 2% paraformaldehyde at 4°C. Cultures were filtered and mounted similar to traditional catalyzed reporter deposition (CARD) FISH methods [41, 42]. To capture choanoflagellate colonies and free-living bacteria, fixed culture was filtered onto a 0.2 µm pore size 25 mm filter (Millipore Sigma, Darmstadt, Germany; Cat. No. GTTP02500). To capture only choanoflagellate colonies and let free-living bacteria pass through, cultures were filtered onto a 5 µm pore size 25 mm filter (Millipore Sigma; Cat. No. TMTP02500). Air-dried filters were coated in 0.1% low melt agarose and cut into wedges for hybridization experiments. To permeabilize bacterial cells, filters were incubated in a CARD-FISH proteinase buffer (10 mg/mL lysozyme; 0.05 M EDTA; 0.1M Tris-HCl, pH 8.0)[42] at 37°C for 30 min. Filters were washed twice in nuclease-free H_2_O, and once in 98% EtOH and left to air-dry. Filters were pre-hybridized in 1 mL of hybridization buffer (100 µl 20X SSC; 100 mg Dextran sulfate (Millipore Sigma; Cat. No. D6001); 200 µl Formamide (20% final conc.)[43] for 30min. at 45°C. Filters were transferred into 500 µl of hybridized buffer with 0.25 µl of 100 mM stock HCR-FISH probes and incubated overnight at 45°C. All filters were labeled with the universal EUB338 probe [44] to label all bacteria. Gam42a was used to label gammaproteobacteria (Fig. 2C’)[45]. Custom probes (Table S4, Table S6) were used to label bacteria from *B. monosierra* cultures. Filters were washed in pre-warmed wash buffer based on the formamide concentration of the hybridization buffer (for 20% formamide hybridization: per 50 mL; 0.5 mL 0.5M EDTA, pH8.0; 1.0 mL 1M Tris HCl, pH8.0; 2150 µl 5M NaCl; 25 µl 20% SDS (w/v))[46]. To wash away unbound probes, filters were incubated in wash buffer for 1 hour at 48°C followed by three washes for five minutes in 5X SSCT (per 40 mL; 10 mL 20X SSC; 400 µl 10% Tween 20). Filters are incubated in amplification buffer (for 40 mL; 10 mL 20X SSC; 8 mL 50% Dextran Sulfate; 400 µl 10% Tween 20)[47] for 30 minutes at room temperature while hairpin solutions were snap cooled as previously described [47]. Signal amplification was performed by incubating filters in amplification buffer with hairpins overnight in the dark in a humidified chamber. To wash, filters were placed in amplification buffer for 1 hour in the dark at room temperature followed by two washes for 30 minutes in 5X SSCT in the dark at room temperature. To stain DNA, filters were washed for 10 minutes in 5X SSCT with 0.1mg/mL Hoechst 33342 (Thermo Fisher Scientific). Finally, filters were washed for 1 minute in nuclease free H_2_O, and 1 minute in 96% EtOH before air drying and mounting in ProLong Diamond (Thermo Fisher Scientific). Slides were left overnight to cure before imaging on a Zeiss Axio Observer LSM 880 with Airyscan detector as described above in confocal imaging. The bacteria SaccML, OceaML1, OceaML2, EctoML1, EctoML2, EctoML3, and RoseML were identified in the culture ML2.1EC. OceaML3, OceaML4, and EctoML4 were identified in the culture ML2.1E.

#### Quantitative image analysis of FISH data set

FISH was performed as described above labeling the microbiome members (SaccML, OceaML1, OceaML2, EctoML1, EctoML2, EctoML3, and RoseML) along with a broad-spectrum bacterial probe EUB 338 [44]. All experiments for quantitative imaging analysis (Fig. S14) were performed in the culture ML2.1EC. A minimum of 120 rosettes were imaged at random per phylotype on 5 µm filters using a 63X/NA 1.15 oil immersion objective on a Zeiss LSM 880 AxioExaminer. A z-stack was acquired for each rosette to capture the whole rosette. Images were analyzed on FIJI [13]. Due to the pressure applied to the rosettes during the filtering process, colonies are flattened; therefore, we applied a maximum projection for each z-stack leaving out slices that contained the filter. Colonies were first assessed to see if the phylotype was present. If at least one bacterial phylotype was found inside the colony, the colony was counted as having the phylotype. Due to the HCR-FISH method, single bacterial resolution is possible (Fig. S12). To determine the abundance of each phylotype present in a colony, colony images were cropped down to only contain the interior microbiome, and then the images were split into separate channels: EUB 338 bacterial probe, phylotype-specific probe, and Hoechst 33342. The Hoechst image was thrown out because it was not needed. An automatic threshold using the Intermodes method [14] was applied to both the broad-spectrum bacterial probe and bacterial phylotype-specific probe images. The area was measured for each threshold (total bacteria broad spectrum probe and phylotype-specific probe) by setting measurements to area and analyzing the images. The area occupied by a particular phylotype was divided by the area of the total bacteria for each colony to determine the proportion or abundance of each bacteria present in individual colonies.

---

**Figure S1:**
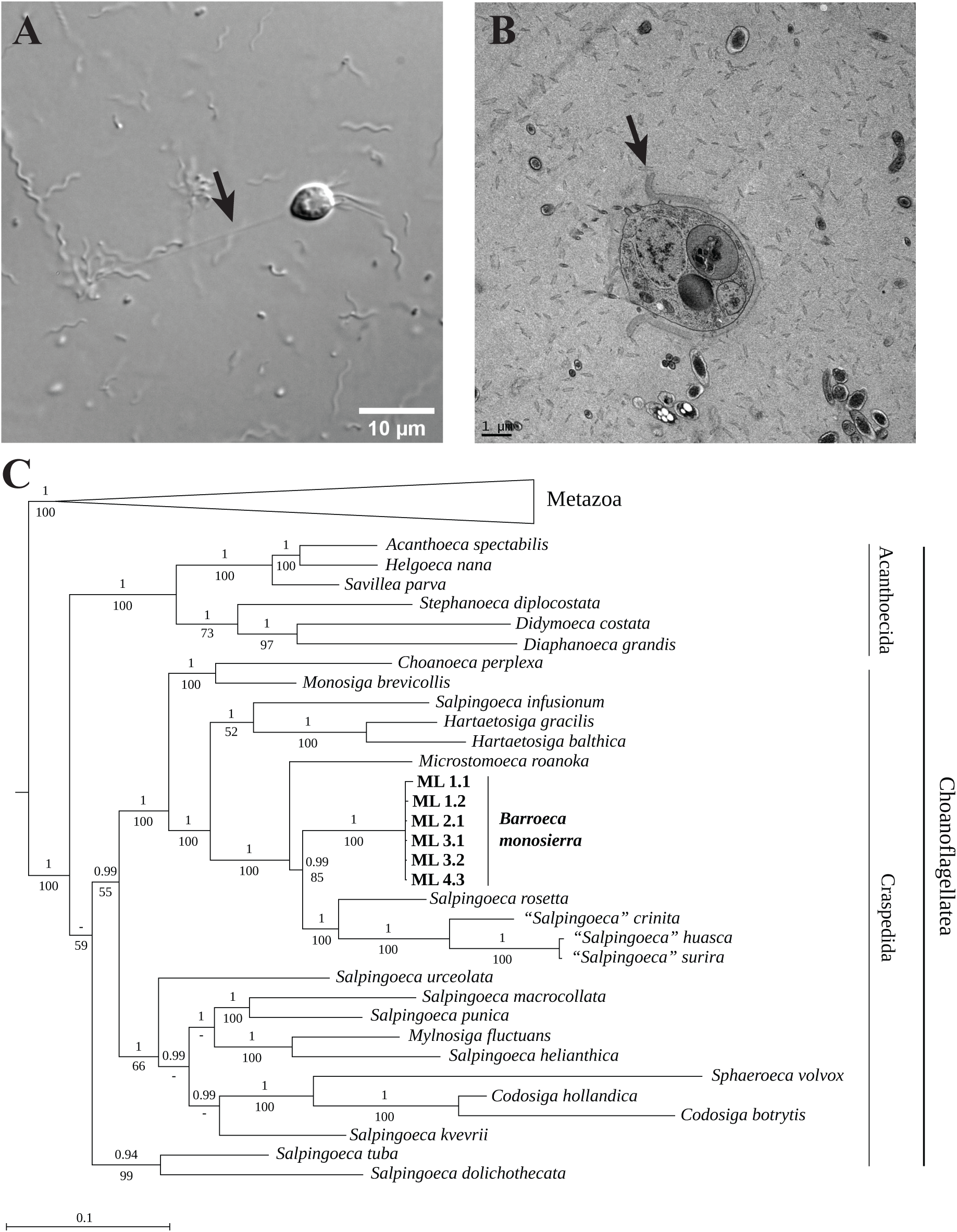
Ultrastructure and phylogeny of choanoflagellate isolates from Mono Lake A. Single cells may be found attached to a substrate via a long (∼30 μm) basal pedicel (arrow). **B**. Single cells may possess an organic cup-shaped theca with a distinctive ∼0.75 μm outward-facing lip on its apical end (arrow). **C**. Phylogenetic tree of sequences from 6 Mono Lake choanoflagellate isolates (labeled on tree) and other choanoflagellate species reveal the isolates (Table S1) are members of the same species. The tree includes data from 3 genes (18S, EFL and HSP90) as a backbone, with each Mono Lake isolate represented only by its 18S sequence. ML 2.1 is the primary strain used in this publication. Metazoa (7 species) were collapsed to save space. Bayesian posterior probabilities are indicated above each internal branch, and maximum likelihood bootstrap values below. (A ‘-’ value indicates a bifurcation lacking support or not present in one of the two reconstructions).

**Figure S2.**
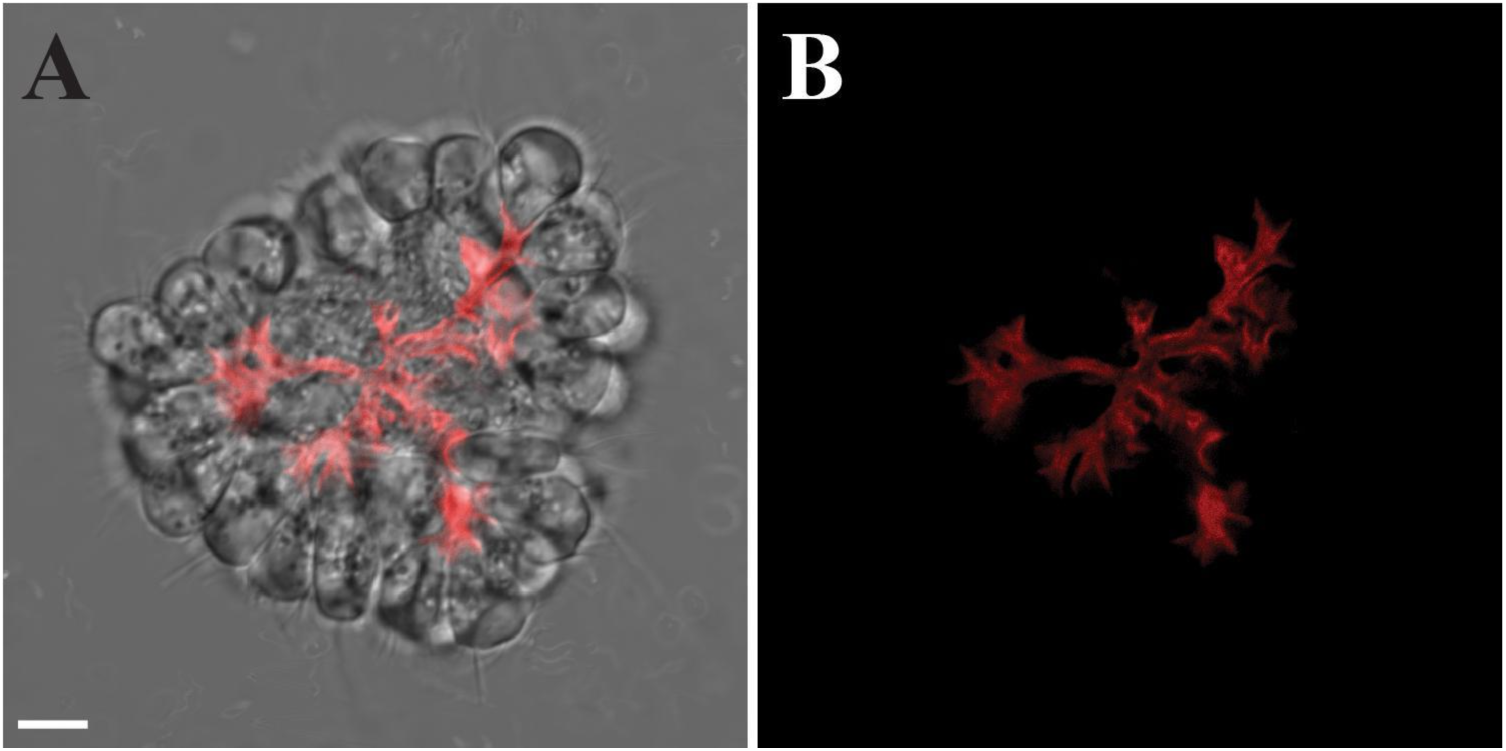
*B. monosierra* colonies contain an extracellular matrix (ECM). (A, B) *B. monosierra* colonies contain a branched network of ECM that extends from and connects the basal pole of cells in the rosette. Optical section of representative colony (A), stained with the lectin Concanavalin A (B; red), shown.

**Figure S3:**
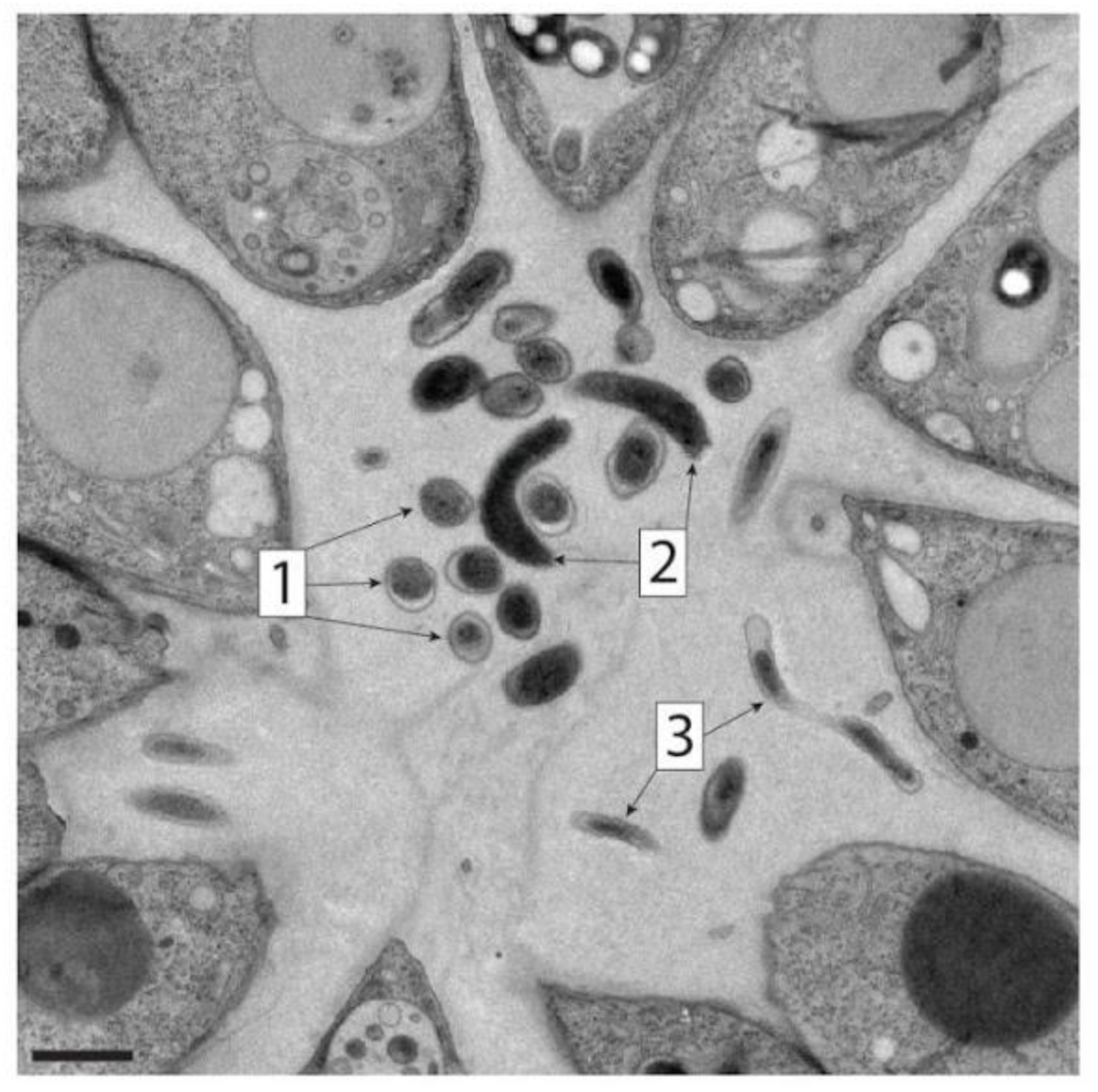
Bacterial residents in *B. monosierra* rosettes exhibit a range of morphologies. Bacteria inside the rosettes of *B. monosierra* have at least 3 distinct morphologies (1-3) revealed by TEM. Scale bar = 1 µm.

**Figure S4:**
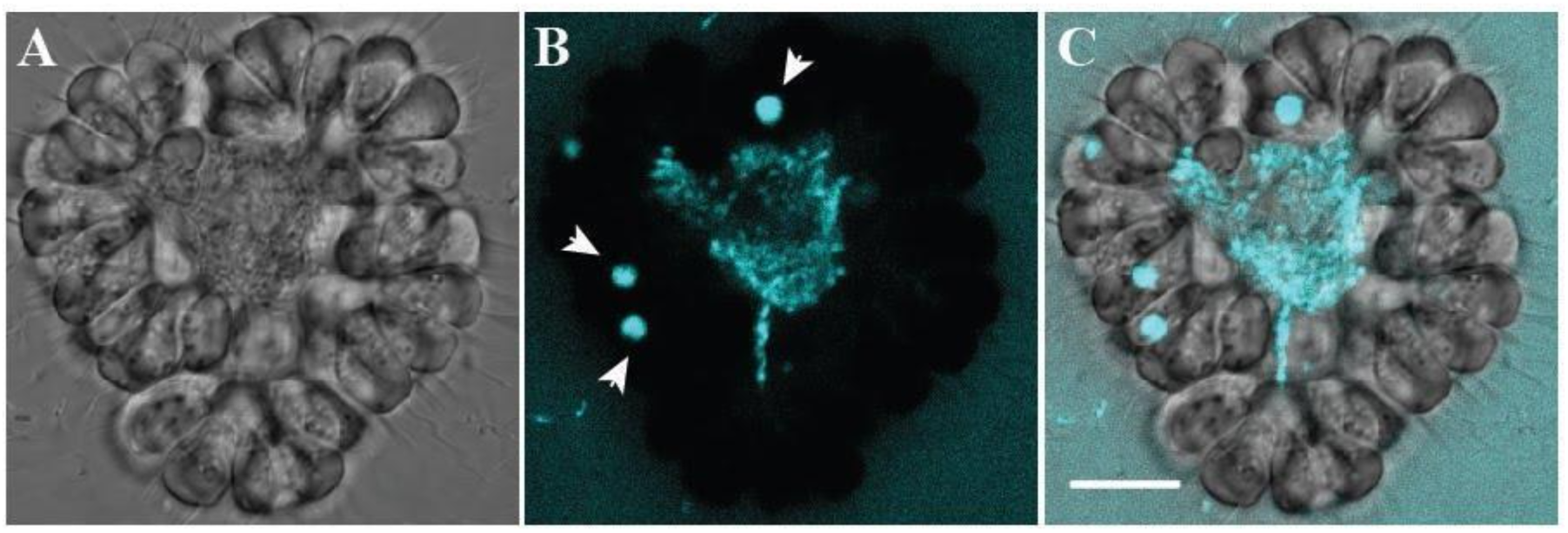
Bacteria inside *B. monosierra* rosettes are alive and growing. Optical section of representative colony (A, DIC). (B) Bacterial cells inside *B. monosierra* colonies incorporate fluorescent D-amino acids into their cell walls (cyan)[15], indicating that they are alive and actively growing. The D-amino acids also accumulate in the food vacuoles of the choanoflagellate cells (arrowheads) through phagocytosis of the dye and of labeled bacteria from outside the colony. Overlay (C) shows D-amino acid enrichment inside the colony, corresponding to the location and morphology of the bacteria in the microbiome. Scale bar = 10 µm.

**Figure S5:**
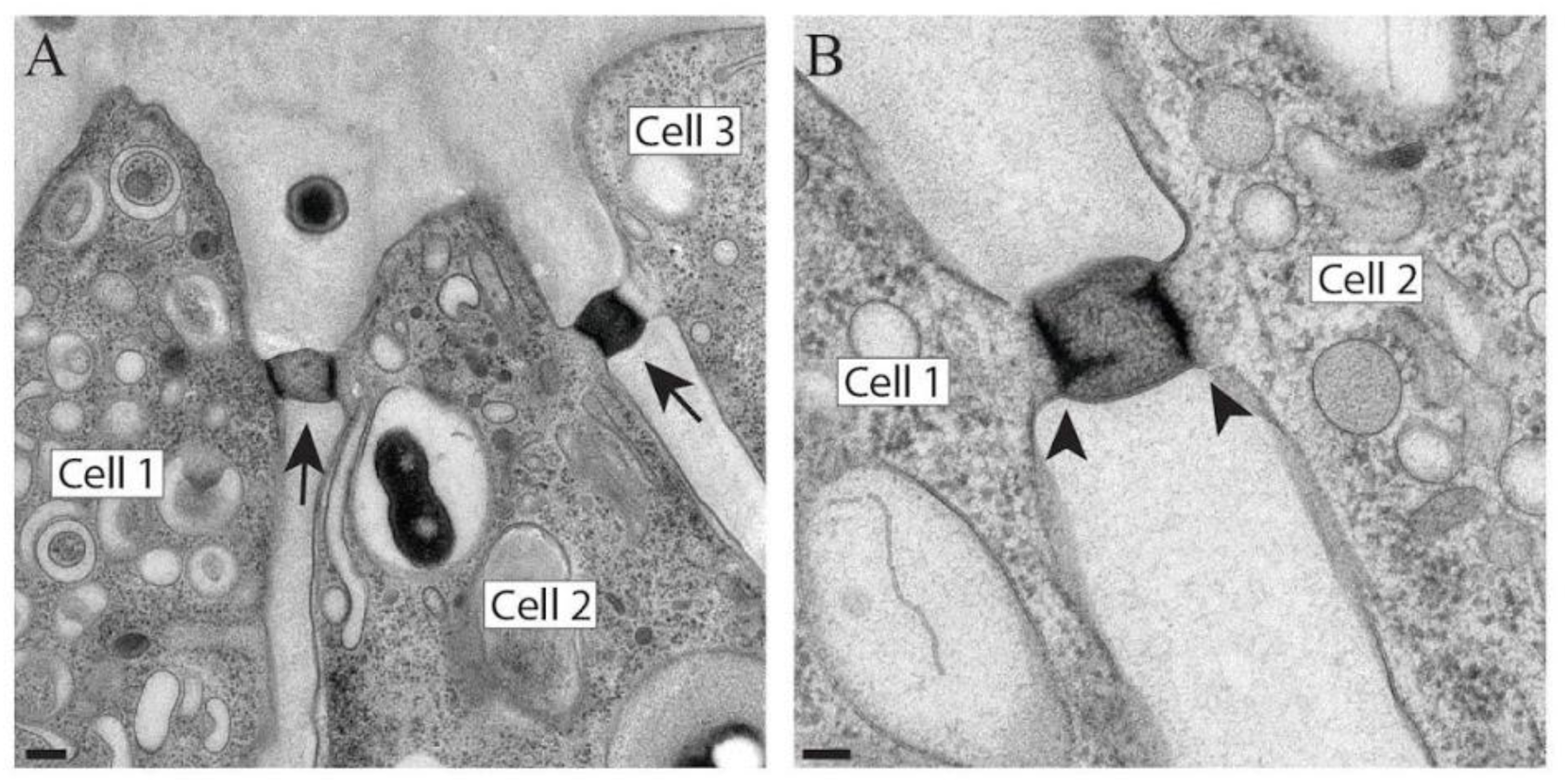
Intercellular bridges connect cells in *B. monosierra* rosettes. (A) Many cells in *B. monosierra* colonies are connected by intercellular bridges (arrows) that resemble the bridges found in *S. rosetta* colonies [16] and other choanoflagellates [48, 49]. Choanoflagellate cells labeled with boxes. (Cell 2 contains a phagocytosed bacterium in its food vacuole. This is not to be confused with the extracellular microbiome illustrated in Figs. 1, 2, and S3.) Scale bar = 200nm. (B) TEM of intercellular bridges between two choanoflagellate cells reveals two electron dense plates (arrowheads) in an arrangement that is reminiscent of the ultrastructure of *S. rosetta* bridges [16]. Scale bar = 100nm.

**Figure S6:**
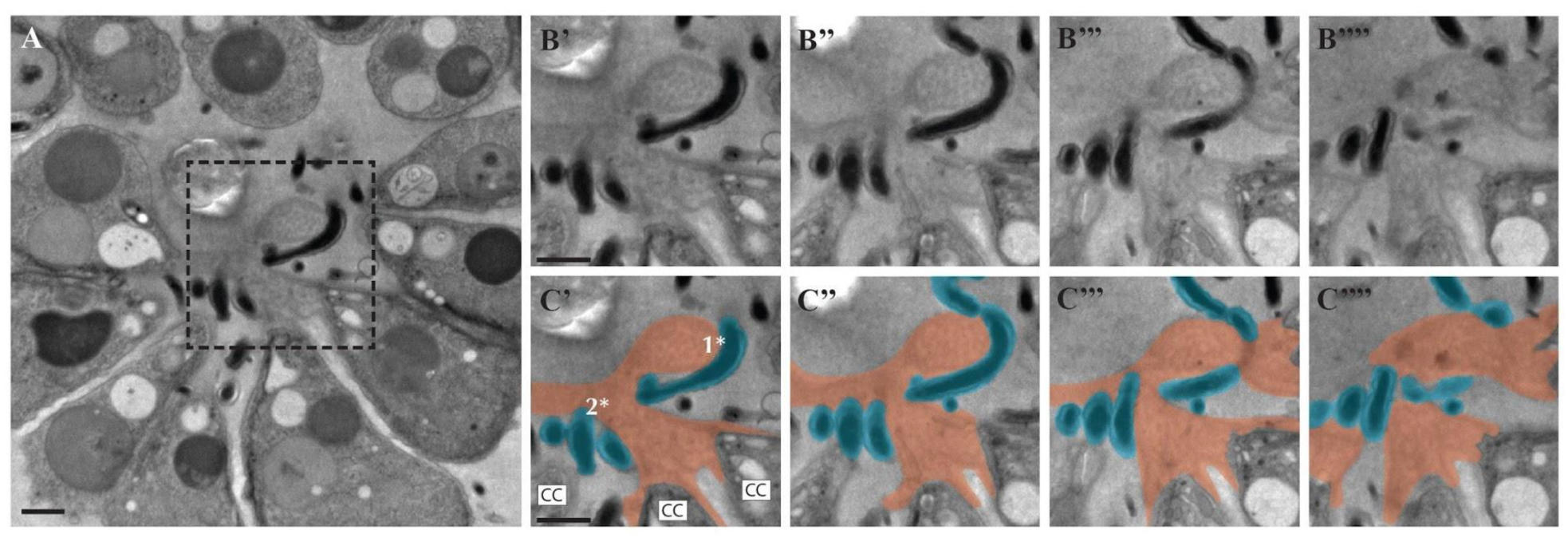
Bacterial residents physically associate with and wrap around the choanoflagellate ECM. (A) A single TEM section through a *B. monosierra* rosette shows choanoflagellate cells on the periphery and bacteria in the center. Box indicates region shown in panels B – C. (B, C) Serial sections (150 nm) through the colony reveal the close proximity of bacteria to the choanoflagellate ECM. (C) False coloring of the TEM sections highlights the associations among the ECM (orange) and bacteria (blue) that wrap around the ECM at two separate sites (1*, 2*). Choanoflagellate cells indicated as CC. Scale bar = 1 µm.

**Figure S7:**
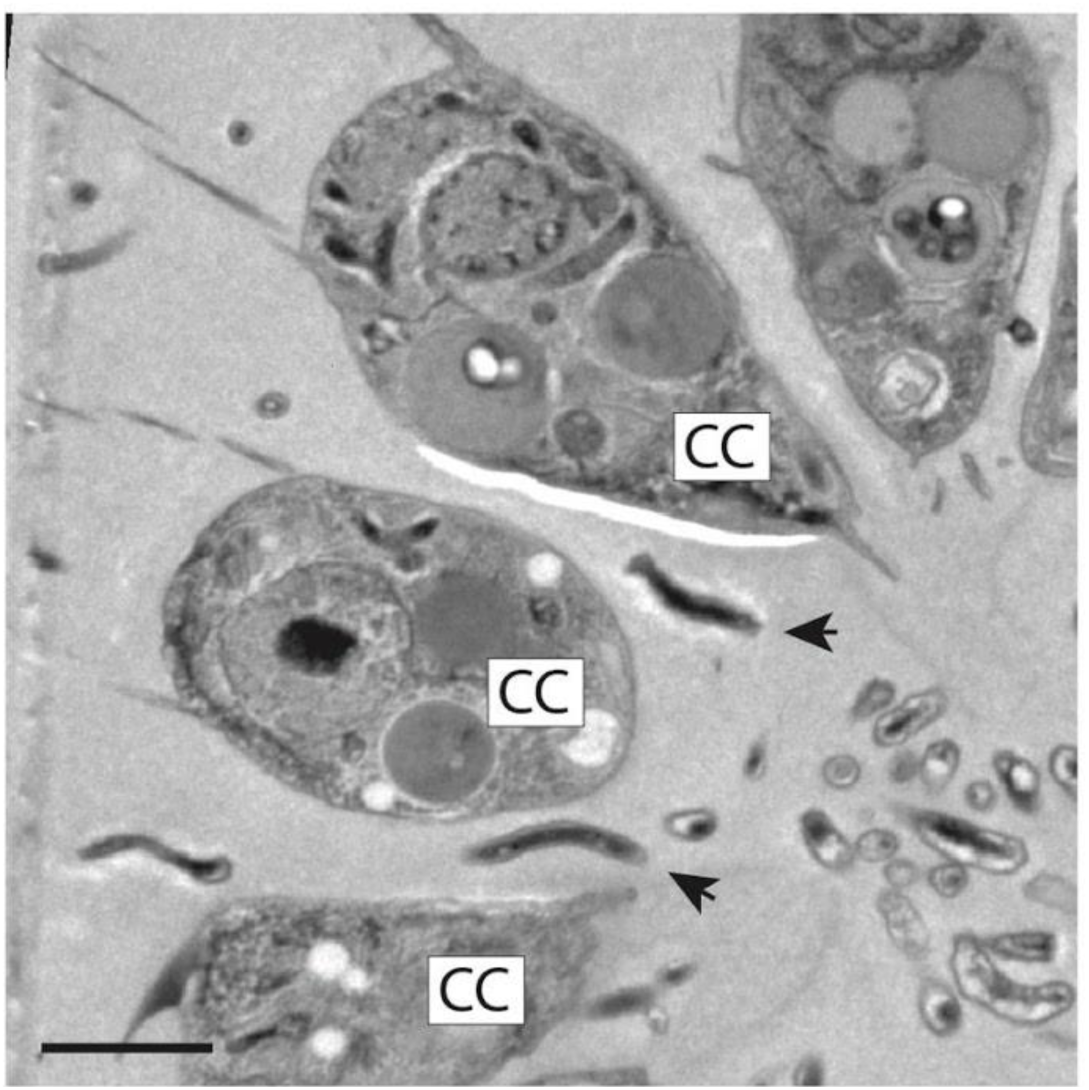
Bacteria are found wedged between the lateral surfaces of *B. monosierra* cells. Bacteria (arrowheads) were observed between cells of *B. monosierra* rosettes. TEM image illustrates two spiral bacteria (arrowheads) positioned between choanoflagellate cells (CC). Scale bar = 2 µm.

**Figure S8:**
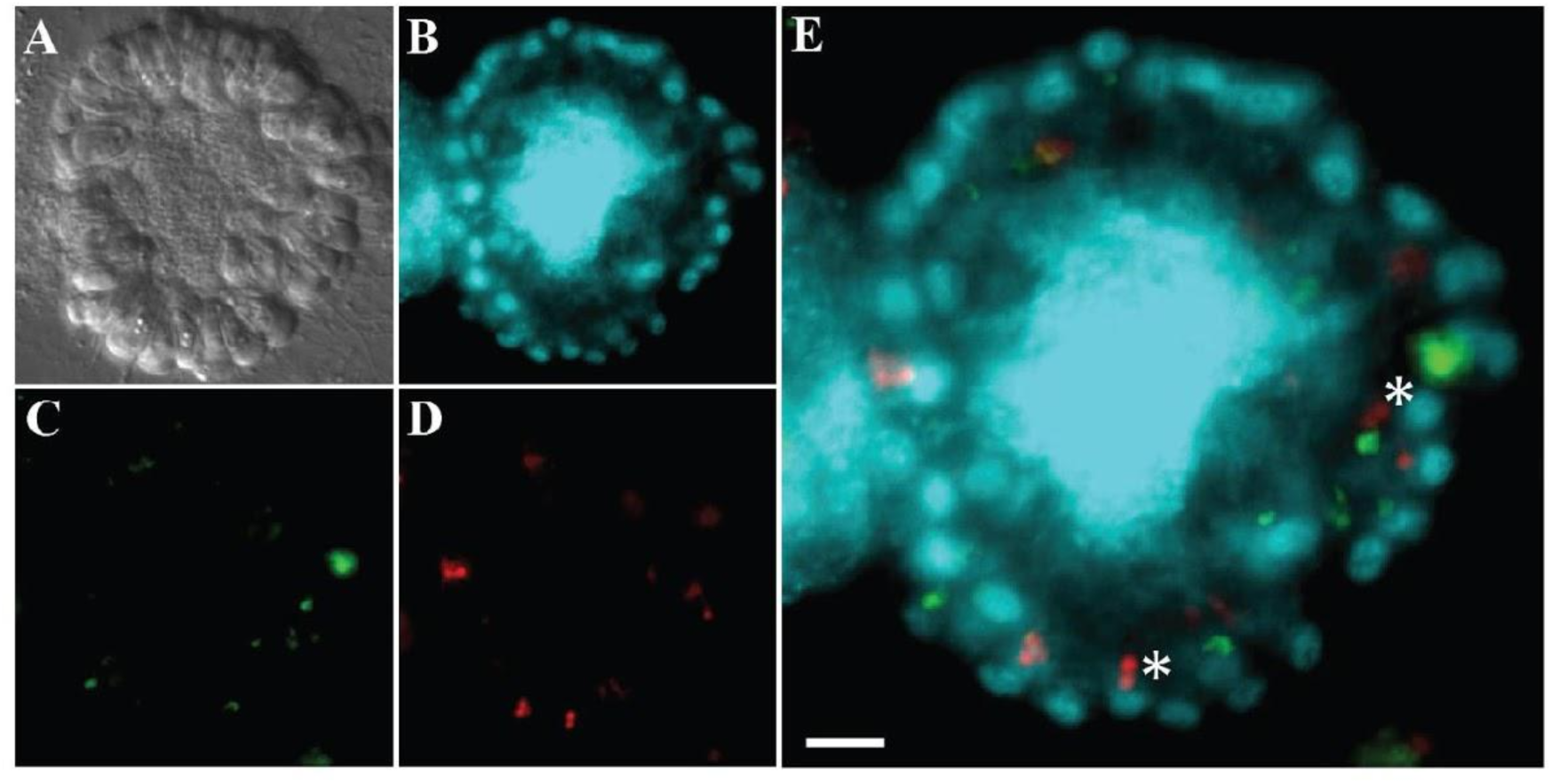
*B. monosierra* rosettes cannot be passively penetrated by sub-micron particles. (A-E) *B. monosierra* colonies fail to incorporate bacteria-sized microspheres over a 24-hour incubation time, suggesting that bacteria cannot enter the colony center passively. An optical section through a representative *B. monosierra* colony (A, DIC; B, Hoechst), illuminates the toroidal choanoflagellate nuclei and interior bacteria. The fluorescent beads (0.22 µm, green, C; 1 µm, red, D) were never observed in the interior cavities of the colonies. They were observed in food vacuoles of choanoflagellate cells (E, asterisks) due to phagocytosis of the beads through the same pathway used for phagocytosis of bacterial prey. Scale bar = 5 µm.

**Figure S9:**
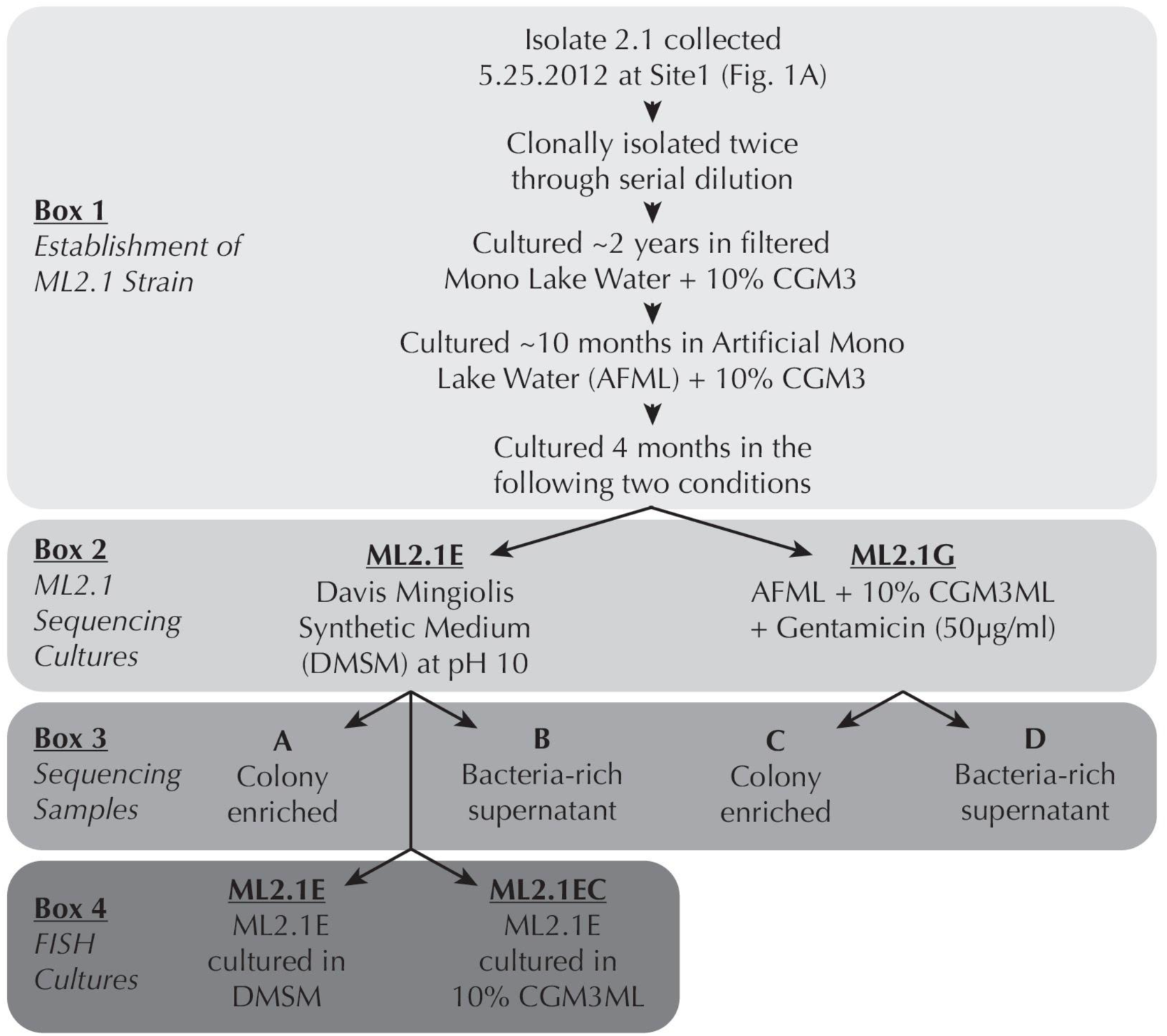
Schematic describing the establishment of *B. monosierra cultures.* ML2.1E and ML2.1G were derived from the original isolate ML2.1 used in the sequencing experiment to identify bacteria present in the *B. monosierra* colonies, as well as, outside of the colonies. (Box 1) Isolate 2.1 was collected on 5.25.2012 at Site 1 (Fig. 1A, Table S1). Each isolate was clonally isolated by serial dilution twice, and ML2.1 was cultured in the lab for nearly 2 years in filtered Mono Lake water supplemented with cereal grass media (CGM3) concentrate as a carbon source. Upon generating an artificial Mono Lake water (AFML), ML2.1 was cultured for 10 months in AFML with CGM3ML. The ML2.1 culture was then split into two cultures (ML2.1E and ML2.1G, Box 2). ML2.1E was cultured in Davis Mingiolis Synthetic minimal medium at pH10 and ML2.1G was similar to ML2.1 cultured in AFML with the addition of a six-week Gentamicin antibiotic treatment that did not affect colony size and improved overall growth of the culture. (Box 3) Through differential centrifugation and filtration, a sample enriched for *B. monosierra* colonies and the supernatant enriched with bacteria were used for genomic DNA extraction for each culture, ML2.1E and ML2.1G. (Box 4) Due to variability in ML2.1E rosette development, likely due to the diversity and complicated growth dynamics of prey bacteria, we found maintaining a culture under multiple growth conditions increased our chances of having a culture with large *B. monosierra* rosettes. The ML 2.1E culture was grown further in DMSM (ML 2.1E) and in 10% CGM3ML (ML 2.1EC); the resulting cultures were used for FISH analysis.

**Figure S10:**
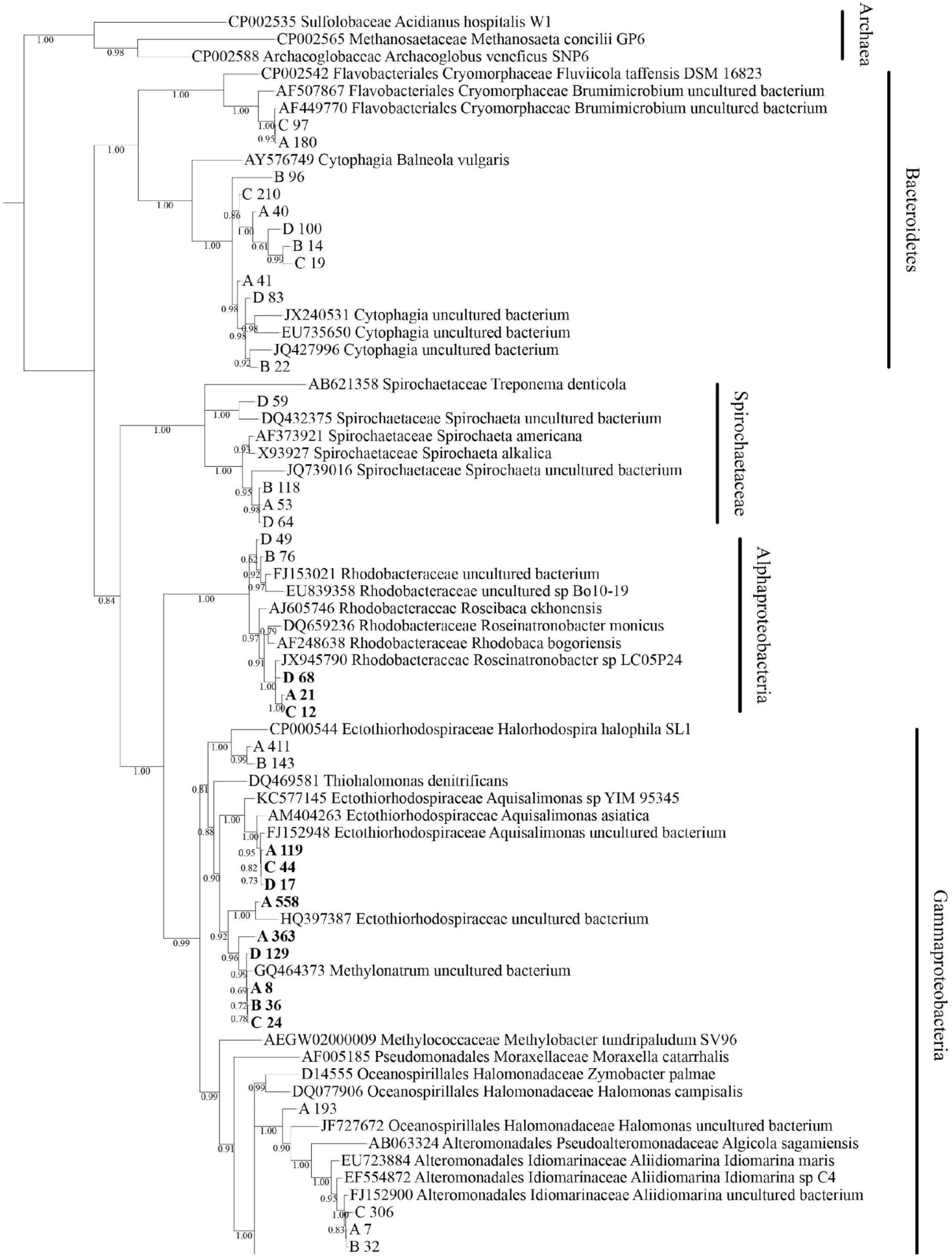
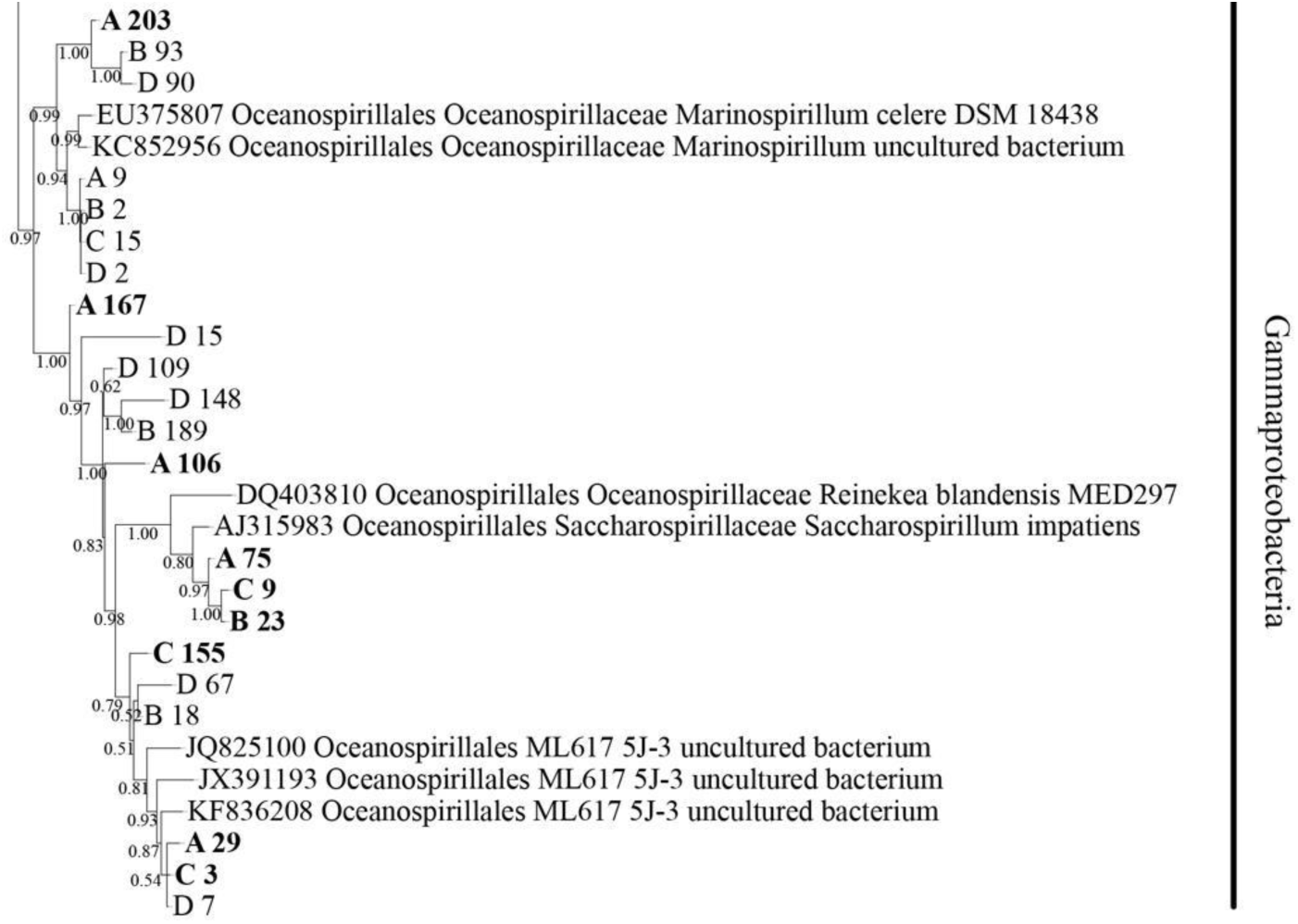
Phylogenetic analysis of 16S rRNA tree of EMIRGE sequences. The tree was built by comparing 16S rRNA sequences assembled from the Mono Lake samples through EMIRGE analysis with 16S rRNA sequences from their closest relatives. Sequences from Mono Lake samples are labeled by the letter associated with the sequencing sample (A or C: with colonies; B or D: environmental bacteria; Fig. S9, Box 3) and a unique identifying number. Species shown in bold were detected inside rosettes (Table S4). Reference species are shown with an accession number and genus name, along with other identifying information. Bootstrap values displayed. Branches with <0.5 bootstrap value have been collapsed.

**Figure S11:**
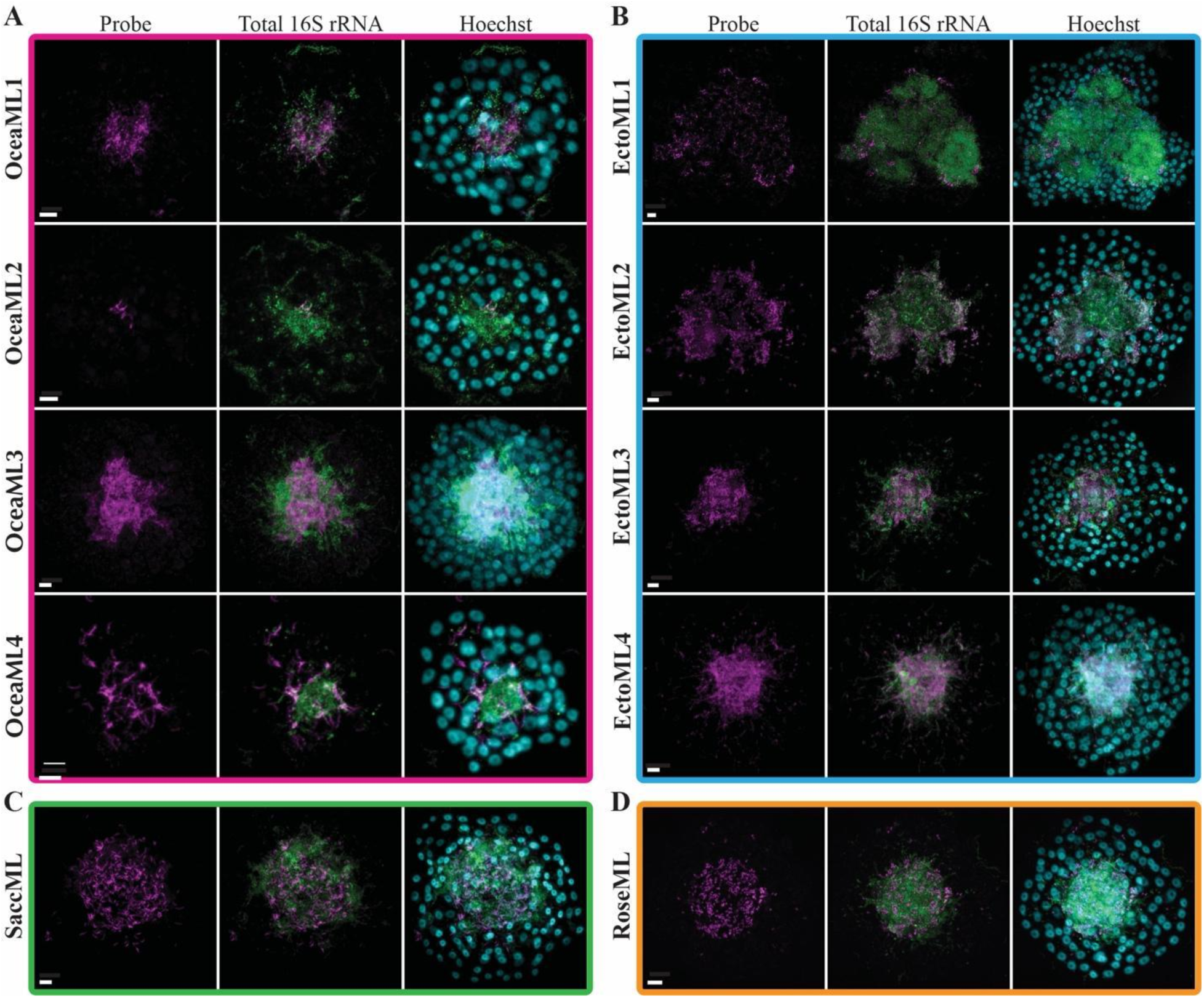
Diverse bacteria are found in the microbiome of *B. monosierra*. (A-D) Ten different bacterial species detected in the center of *B. monosierra* colonies. Confocal images of representative colonies hybridized with phylotype-specific probes (magenta; Table S4, Table S6) and a broad spectrum 16S rRNA probe (green) overlaid with Hoechst 33342 staining of the choanoflagellate nuclei and bacterial nucleoids (cyan). The bacteria are grouped by genus and color coded to match the phylogenetic tree in Fig. 2E. The bacteria have been named as follows: (A) OceaML1 = *Oceanospirillaceae* sp.1; OceaML2 = *Oceanospirillaceae* sp. 2; OceaML3 = *Oceanospirillaceae* sp.3; OceaML4 = *Oceanospirillaceae* sp. 4; (B) SaccML = *Saccharospirillaceae* sp.; (C) EctoML1 = *Ectothiorhodospiraceae* sp. 1; EctoML2 = *Ectothiorhodospiraceae* sp. 2; EctoML3 = *Ectothiorhodospiraceae* sp. 3; EctoML4 = *Ectothiorhodospiraceae* sp. 4. and (D) RoseML = *Roseinatronobacter* sp. Scale bars = 5 µm.

**Figure S12:**
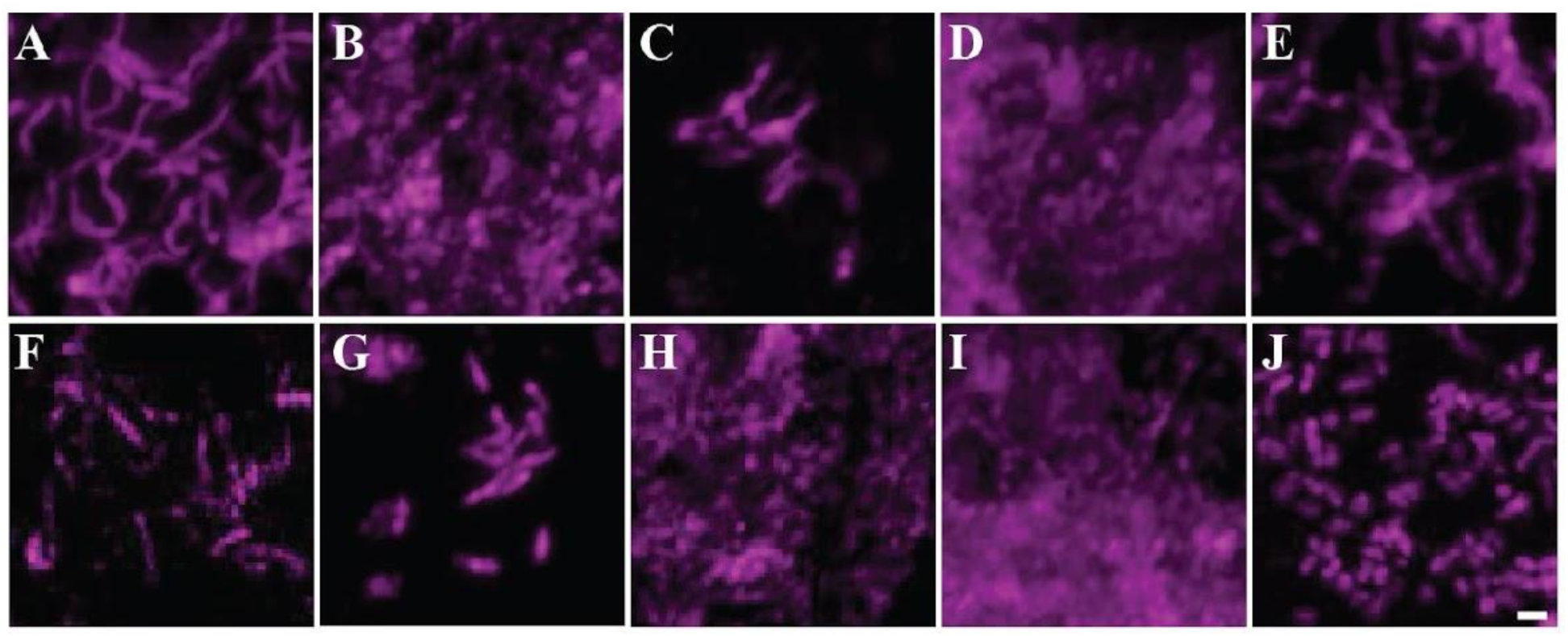
*B. monosierra* microbiota members exhibit filamentous and rod morphologies. HCR-FISH probes illuminate distinct bacterial morphologies, from filamentous bacteria (A, C, E, F) to rod-shaped bacteria (B, G, J). (A) SaccML, (B) OceaML1, (C) OceaML2, (D) OceaML3, (E) OceaML4, (F) EctoML1, (G) EctoML2, (H) EctoML3, (I) EctoML4, (J) RoseML. Scale bar = 1 µm.

**Figure S13:**
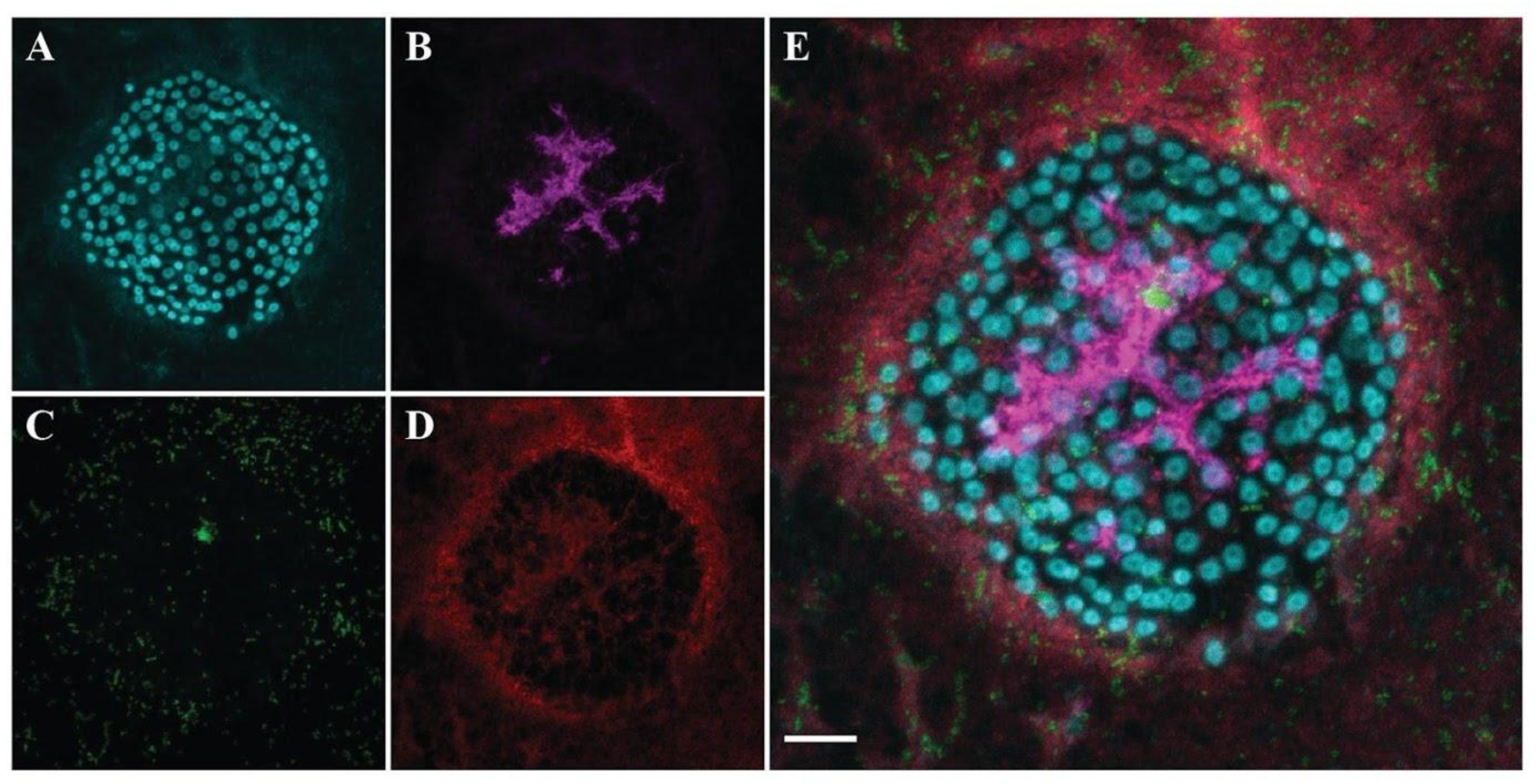
The bacterium OceaML3 was detected exclusively inside rosettes. The *B. monosierra* colony identified by the cluster of toroidal nuclei (A, cyan), shows how OceaML3 (B, magenta) is present exclusively inside the rosette whereas EctoML2 (C, green) and EctoML4 (D, red) can be detected both inside and outside the rosette. (E) Merge of all four channels. FISH analysis was performed on a 0.22 µm filter to capture free-living bacteria. Scale bar = 10 µm.

**Figure S14:**
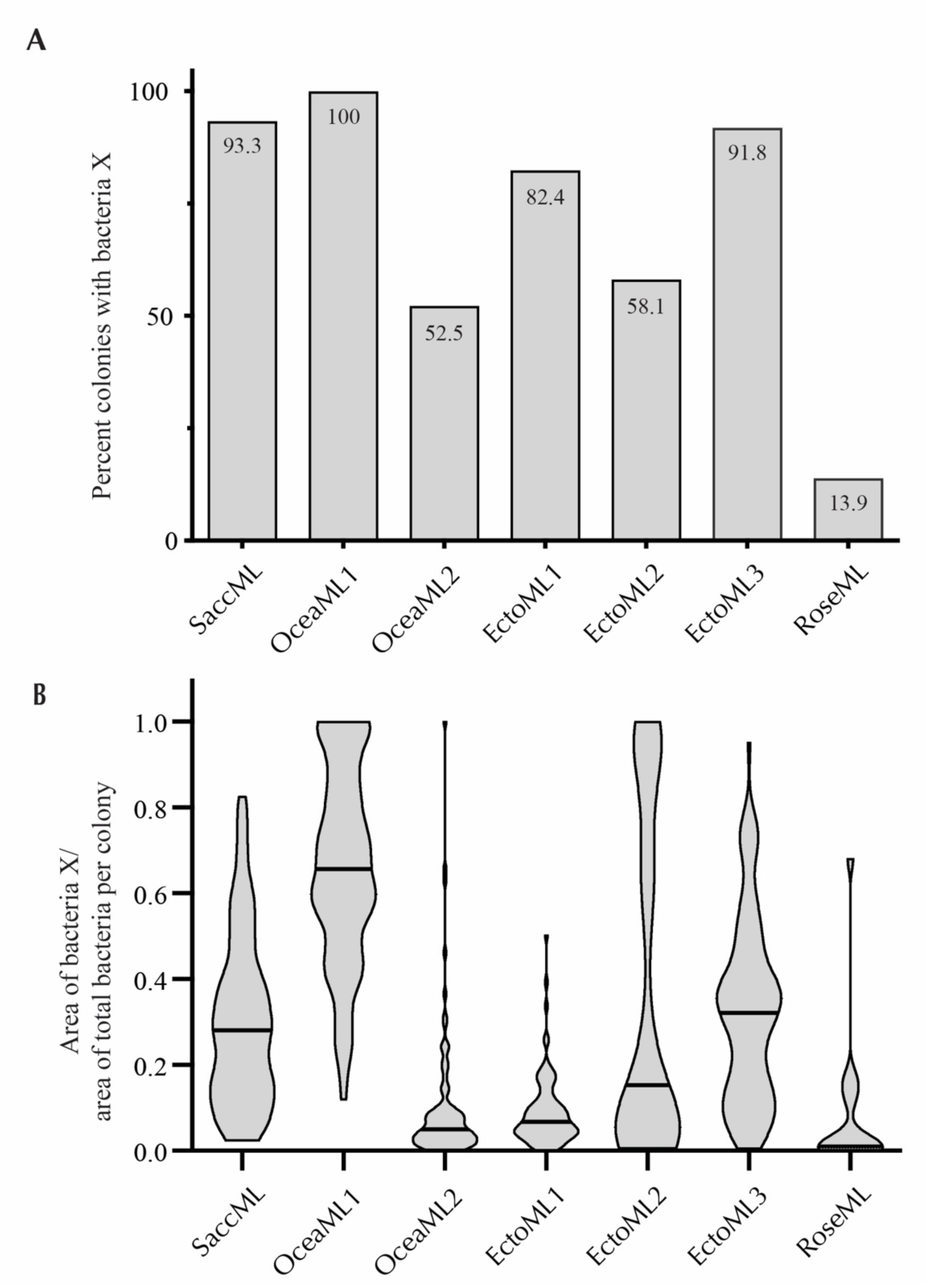
OceaML1 is a core member of the *B. monosierra* microbiome. Frequency of bacteria (A) and their relative abundance inside the rosette (B) illustrates that each bacterium has a unique pattern. (A) The presence of each phylotype was assessed for a minimum of 120 rosettes. HCR-FISH can resolve individual bacteria, and only one bacterium was needed to confirm presence. (B) The area of the bacteria compared to the total area was used to determine the relative abundance. The data were graphed as a violin plot with the mean indicated as the black line.

**Figure S15:**
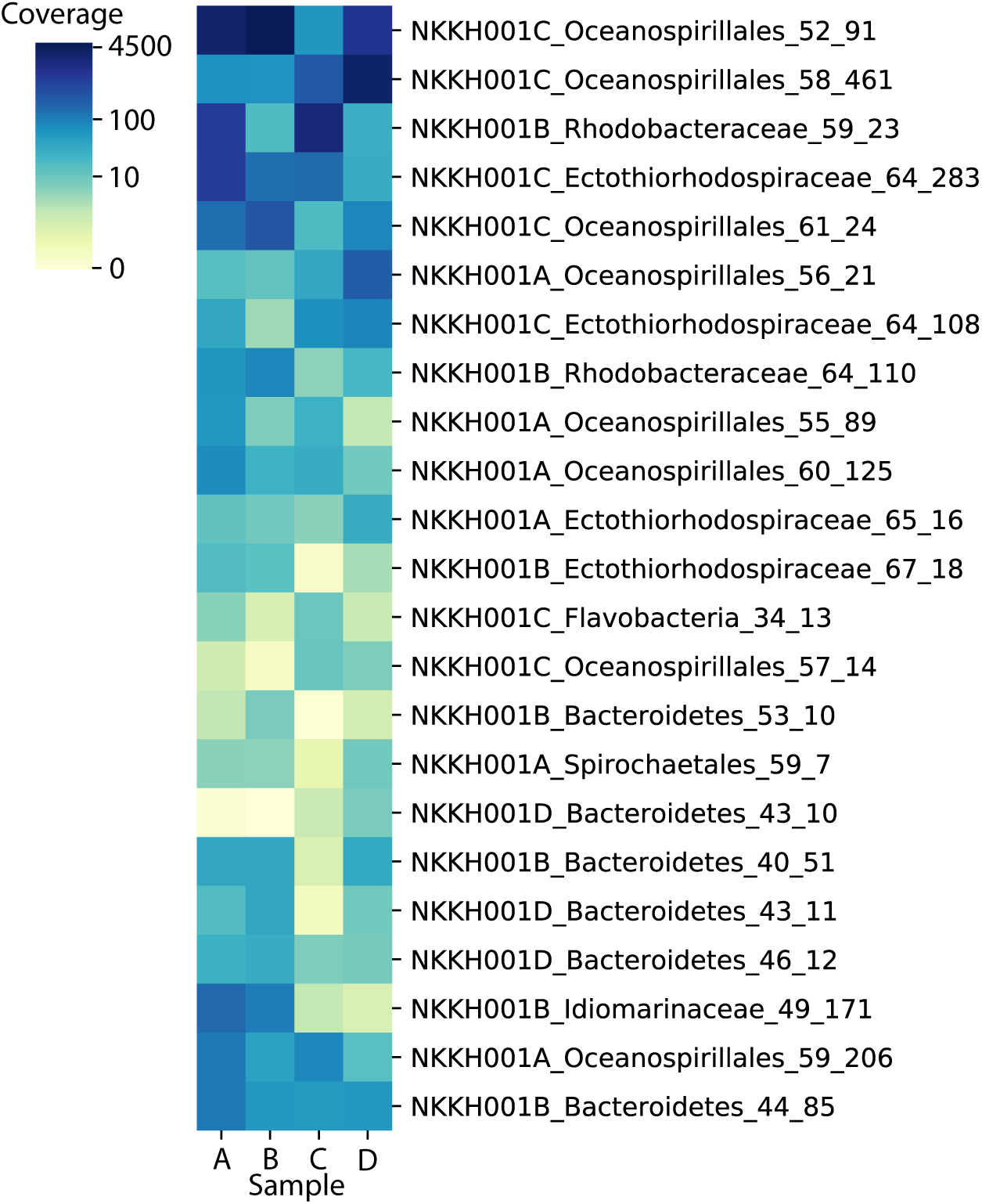
Bacterial community overlap across shotgun metagenomic sequencing samples. Sequencing reads from each shotgun metagenome were cross-mapped against the dereplicated bin set. Coverage for each bin was determined by averaging the coverage of a given bin’s constituent contigs.

**Table S1:** Date, location, phenotype, and isolate designations for *Barroeca monosierra* isolates.

| **Isolate** | **Date collected** | **Location\*\*** | **18S Sequenced** | **% Identity with ML2.1** | **Phenotype when isolated** | **Phenotype after culturing in lab** |
| --- | --- | --- | --- | --- | --- | --- |
| ML 1.1 | 05.15.2012 | Site 1 | Yes | 99.6% | Single Celled | Colonies |
| ML 1.2 | 05.15.2012 | Site 1 | Yes | 99.4% | Single Celled | Colonies |
| ML 2.1* | 05.25.2012 | Site 1 | Yes | 100% | Colonies | Colonies |
| ML 3.1 | 05.05.2013 | Site 1 | Yes | 99.3% | Colonies | Colonies |
| ML 3.2 | 05.05.2013 | Site 1 | Yes | 99.3% | Colonies | Colonies |
| ML 3.3 | 05.05.2013 | Site 1 | No |  | Colonies | Colonies |
| ML 4.1 | 10.11.2014 | Site 2 | No |  | Colonies | Colonies |
| ML 4.2 | 10.11.2014 | Site 2 | No |  | Colonies | Colonies |
| ML 4.3 | 10.11.2014 | Site 2 | Yes | 99.5% | Colonies | Colonies |
| ML 4.4 | 10.11.2014 | Site 2 | No |  | Colonies | Colonies |

\* Primary strain used in this publication
\*\* See Fig. 1A. Site 1 is at the Mono Lake picnic area approximately at 37°58'42.7"N 119°01'52.9"W. Site 2 is at the Mono Lake South Tufa area approximately at 37°56'37.1"N 119°01'37.8"W.

**Table S2:** Shotgun metagenomic sequencing project outcomes

|  | Sample |  |  |  |
| --- | --- | --- | --- | --- |
|  | A | B | C | D |
| Sequencing Depth (Gbp) | 34.1 | 25.8 | 31.3 | 22.6 |
| Assembly size (Mbp) | 173.7 | 86.7 | 141.6 | 87.4 |
| EukRep removed sequence (Mbp) | 50.3 | NA | 51.7 | NA |
| # bins | 17 | 12 | 9 | 10 |
| # bins (dereplicated) | 6 | 7 | 7 | 3 |

**Table S3:** Nine genera of bacteria identified in *B. monosierra* cultures through two independent analyses: comparison of ribosomal proteins detected through metagenomic assembly and by 16s rRNA assembly and analysis.

| Class | EukRep Metagenomic Analysis* |  | EMIRGE 16S rRNA Analysis* |  |
| --- | --- | --- | --- | --- |
|  | Genus | Number of species | Genus | Number of species |
| *Spirochaetia* | *Spirochaetia sp.* | 2 | *Spirochaetia sp.* | 2 |
| *Gammaproteobacteria* | *Oceanospirillaceae sp.* | 6 | *Oceanospirillaceae sp.* | 6 |
|  | *Saccharospirillaceae sp.* | 1 | *Saccharospirillaceae sp.* | 1 |
|  | *Ectothiorhodospiraceae sp.* | 4 | *Ectothiorhodospiraceae sp.* | 5 |
|  | *Idiomarinaceae sp.* | 1 | *Idiomarinaceae sp.* | 1 |
|  | *Marinospirillum sp.* | 1 | *Marinospirillum sp.* | 2 |
| *Alphaproteobacteria* | *Rhodobacteraceae sp.* | 2 | *Rhodobacteraceae sp.* | 2 |
| *Bacteroidetes* | *Chitonophagaceae sp.* | 6 | *Chitonophagaceae sp.* | 4* |
|  | *Fluviicola sp.* | 1 | *Fluviicola sp.* | 1 |
|  | Total | 24 | Total | 24 |

\*See methods for detailed explanation of the relative challenges in using either solely 16S rRNA analysis or metagenomic analysis to detect uncharacterized bacteria from environmental samples.

**Table S4:** Predicted targets of HCR-FISH probes based on 16S rRNA sequences. Bacteria in bold were identified inside *B. monosierra* colonies.

| Bacteria * | Probe name | Target 16S EMIRGE Sequence** |
| --- | --- | --- |
| ***Saccharospirillum* sp.** | KH 133_Sacc3 | ML_16S_A_75,<br>ML_16S_B_23,<br>ML_16S_C_9 |
| *Idiomarinaceae* sp. | OG_Idio1 | ML_16S_A_7,<br>ML_16S_B_32,<br>ML_16S_C_306 |
| *Marinospirillum* sp.1 | OG_Mar1 | ML_16S_A_193 |
| *Marinospirillum* sp.2 | OG_Mar1 | ML_16S_A_9,<br>ML_16S_B_2,<br>ML_16S_C_15,<br>ML_16S_D_2 |
| ***Oceanospirillales* sp.1** | KH 270_OceaA167.3 | ML_16S_A_167 |
| ***Oceanospirillales* sp.2** | KH 278_OceaA29.1 | ML_16S_A_29 |
| ***Oceanospirillales* sp.3** | KH 295_OceaC3.5 | ML_16S_C_3 |
| ***Oceanospirillales* sp.4-5\*\*** | KH 288_OceaA106_C155 | ML_16S_A_106,<br>ML_16S_C_155 |
| ***Oceanospirillales* sp.5-6\*\*** | KH 293_OceaA203_C155.1 | ML_16S_A_203,<br>ML_16S_C_155 |
| ***Ectothiorhodospiraceae* sp.1** | KH 168_Ecto8 | ML_16S_A_363 |
| ***Ectothiorhodospiraceae* sp.2** | KH 171_Ecto11 | ML_16S_A_8,<br>ML_16S_B_36,<br>ML_16S_C_24,<br>ML_16S_D_129 |
| ***Ectothiorhodospiraceae* sp.3** | KH 174_Ecto14 | ML_16S_A_119,<br>ML_16S_C_44,<br>ML_16S_D_17 |
| ***Ectothiorhodospiraceae* sp.4** | KH 297_EctoA558.4 | ML_16S_A_558 |
| *Ectothiorhodospiraceae* sp.5 | KH 285_EctoA411.3 | ML_16S_A_411,<br>ML_16S_B_143 |
| ***Roseinatronobacter* sp.** | KH 152_Rose3 | ML_16S_A_21,<br>ML_16S_C_12,<br>ML_16S_D_68 |
| *Rhodobacterales* sp. | KH 154_Rhod2 | ML_16S_B_76,<br>ML_16S_D_49 |
| *Cytophagia* sp.1 | *** | ML_16S_B_96 |
| *Cytophagia* sp.2 | *** | ML_16S_B_22,<br>ML_16S_D_83 |
| *Cytophagia* sp.3-4 **** | KH 177_Cyto2 | ML_16S_A_40,<br>ML_16S_A_41,<br>ML_16S_B_14,<br>ML_16S_C_19,<br>ML_16S_C_210,<br>ML_16S_D_100 |
| *Brumimicrobium* sp. | KH 124_Brum3 | ML_16S_A_180,<br>ML_16S_C_97 |
| *Spirochaetia* sp. | KH 148_Spir2 | ML_16S_A_53,<br>ML_16S_B_118,<br>ML_16S_D_64 |

\*\* Target 16S rRNA sequences from EMIRGE analysis. A and C sequences are from *B. monosiera* colony-enriched samples while B and D sequences are from the bacteria-rich supernatant.
\*\*\* Bacteria identified only in bacteria-rich supernatant and not in colony-enriched sequences.
\*\*\*\* Exact number of species is undetermined due to sequence similarity in 16S rRNA EMIRGE assembly from 150bp Illumina sequencing reads.

**Table S5:** Bacterial 16S rRNA – Genbank accession numbers

| **#Accession** | **Sequence ID** |
| --- | --- |
| MW827060 | A_9 |
| MW827061 | A_21 |
| MW827062 | A_29 |
| MW827063 | A_167 |
| MW827064 | A_7 |
| MW827065 | A_203 |
| MW827066 | A_8 |
| MW827067 | A_75 |
| MW827068 | A_106 |
| MW827069 | A_40 |
| MW827070 | A_41 |
| MW827071 | A_119 |
| MW827072 | A_411 |
| MW827073 | A_53 |
| MW827074 | A_558 |
| MW827075 | A_180 |
| MW827076 | B_2 |
| MW827077 | B_18 |
| MW827078 | B_189 |
| MW827079 | B_32 |
| MW827080 | B_76 |
| MW827081 | B_36 |
| MW827082 | B_22 |
| MW827083 | B_14 |
| MW827084 | B_96 |
| MW827085 | B_143 |
| MW827086 | B_23 |
| MW827087 | B_118 |
| MW827088 | C_12 |
| MW827089 | C_3 |
| MW827090 | C_155 |
| MW827091 | C_15 |
| MW827092 | C_9 |
| MW827093 | C_24 |
| MW827094 | C_44 |
| MW827095 | C_19 |
| MW827096 | C_210 |
| MW827097 | C_306 |
| MW827098 | C_97 |
| MW827099 | D_7 |
| MW827100 | D_2 |
| MW827101 | D_109 |
| MW827102 | D_67 |
| MW827103 | D_15 |
| MW827104 | D_148 |
| MW827105 | D_17 |
| MW827106 | D_100 |
| MW827107 | D_49 |
| MW827108 | D_68 |
| MW827109 | D_129 |
| MW827110 | D_83 |
| MW827111 | D_59 |
| MW827112 | D_64 |

**Table S6:**
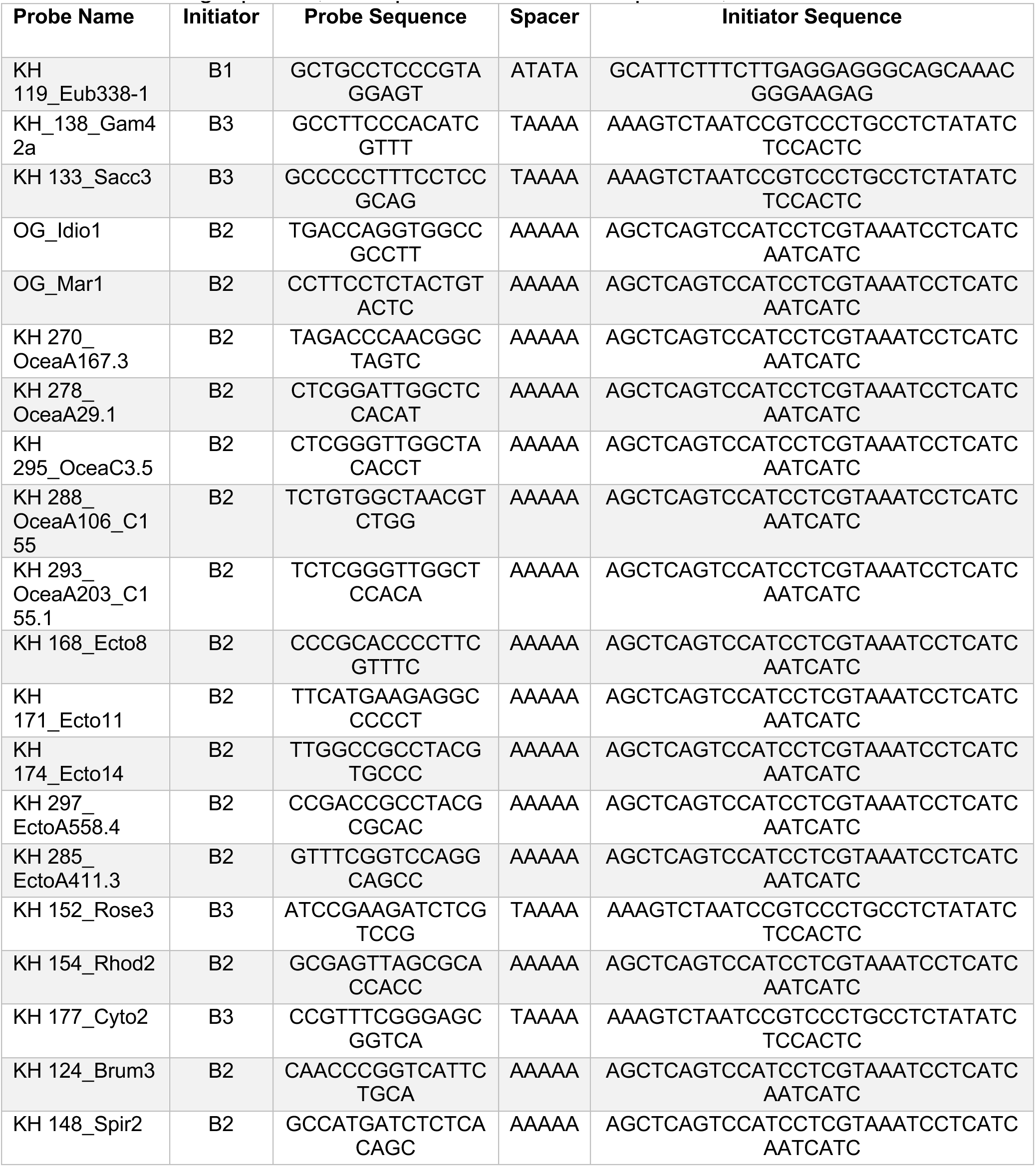
Full length probes, with spacer and initiator sequences, used for HCR-FISH.

**Table S7:**
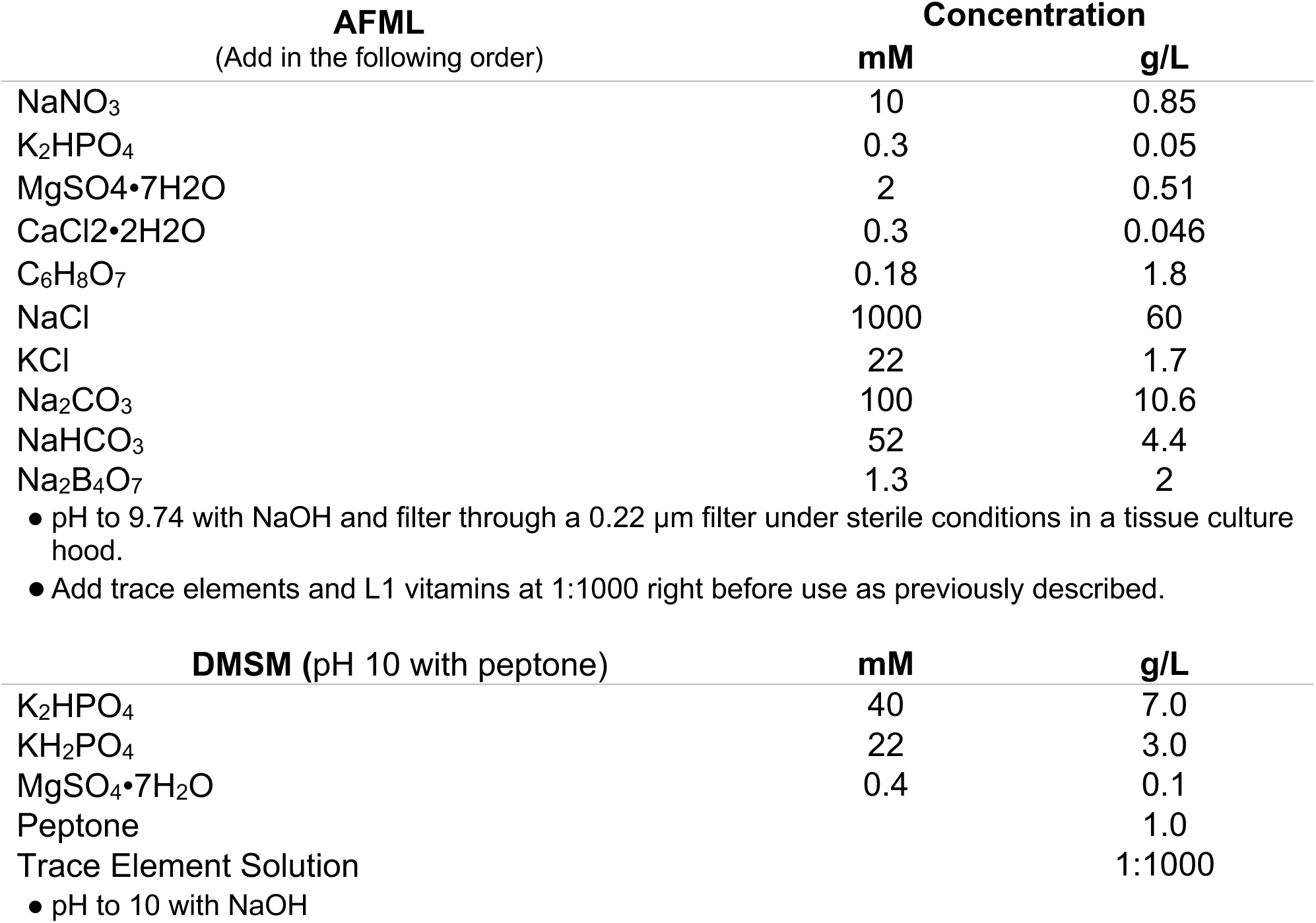
**Growth Media Recipes** [6, 8][50]

**Table S8:** Dereplicated bin set used for constructing the ribosomal protein tree

| **Bin** | **Size (Mbp)** | **Coverage** | **Completeness** |
| --- | --- | --- | --- |
| NKKH001A_Ectothiorhodospiraceae_65_16 | 2.86 | 15.96 | 0.91 |
| NKKH001A_Oceanospirillales_55_89 | 3.74 | 88.50 | 0.95 |
| NKKH001A_Oceanospirillales_56_21 | 4.17 | 21.26 | 0.85 |
| NKKH001A_Oceanospirillales_59_206 | 3.08 | 206.07 | 0.95 |
| NKKH001A_Oceanospirillales_60_125 | 4.32 | 124.61 | 0.93 |
| NKKH001A_Spirochaetales_59_7 | 1.72 | 7.09 | 0.89 |
| NKKH001B_Bacteroidetes_40_51 | 4.13 | 50.72 | 0.91 |
| NKKH001B_Bacteroidetes_44_85 | 3.44 | 84.76 | 0.95 |
| NKKH001B_Bacteroidetes_53_10 | 3.45 | 9.58 | 0.87 |
| NKKH001B_Ectothiorhodospiraceae_67_18 | 2.45 | 18.41 | 0.93 |
| NKKH001B_Idiomarinaceae_49_171 | 2.82 | 170.77 | 0.93 |
| NKKH001B_Rhodobacteraceae_59_23 | 4.00 | 23.36 | 0.95 |
| NKKH001B_Rhodobacteraceae_64_110 | 4.66 | 113.54 | 0.95 |
| NKKH001C_Ectothiorhodospiraceae_64_108 | 3.58 | 108.39 | 0.95 |
| NKKH001C_Ectothiorhodospiraceae_64_283 | 2.92 | 283.97 | 0.95 |
| NKKH001C_Flavobacteria_34_13 | 3.42 | 13.12 | 0.95 |
| NKKH001C_Oceanospirillales_52_91 | 3.42 | 91.11 | 0.93 |
| NKKH001C_Oceanospirillales_57_14 | 3.58 | 14.03 | 0.85 |
| NKKH001C_Oceanospirillales_58_461 | 3.56 | 459.08 | 0.95 |
| NKKH001C_Oceanospirillales_61_24 | 3.17 | 24.31 | 0.93 |
| NKKH001D_Bacteroidetes_43_10 | 3.66 | 10.19 | 0.91 |
| NKKH001D_Bacteroidetes_43_11 | 3.49 | 11.21 | 0.93 |
| NKKH001D_Bacteroidetes_46_12 | 4.54 | 11.51 | 0.91 |

